# Conservation rescued the Mauritius kestrel from extinction but not from genomic erosion

**DOI:** 10.64898/2026.06.23.733933

**Authors:** Xuejing Wang, Anna Stuart, Ken Norris, Sion Henshaw, Alexandra Strang, Carolina Pacheco, Sascha Dreyer Nielsen, Madeleine Waite, Julian P. Hume, Kevin Ruhomaun, Selina Brace, M. Thomas P. Gilbert, Vikash Tatayah, Carl Jones, Jim Groombridge, Cock van Oosterhout, Hernán E. Morales

## Abstract

Conservation can prevent species extinction via demographic recovery, yet it remains debated whether this translates into genomic recovery and restored fitness. The Mauritius kestrel (*Falco punctatus*) declined to four known wild birds in 1974 before intensive management recovered the population. Using 130 genomes spanning nearly 200 years, lifetime reproductive success data, and simulations, we reconstructed genomic change across the species’ collapse and recovery. Long-term small population size had already removed some harmful variation before the crash, a process expected to buffer populations from severe inbreeding depression. Yet the recent bottleneck sharply increased inbreeding, exposed additional harmful variants, and left a signature of genomic erosion associated with reduced reproductive success. The long conservation history of the Mauritius kestrel shows how population collapse and recovery can leave a compounding genetic threat, in which partial genetic purging, continuing genomic erosion, and conservation dependence unfold together in rescued species.

## Introduction

Many species now persist as small, fragmented, or recently rescued populations, raising a central question for conservation: does demographic recovery restore long-term evolutionary viability? Conservation action has averted the extinction of numerous species^1–3^, e.g. birds^4–7^ and mammals^8–10^, demonstrating that population numbers can recover even after extreme decline^11^. However, population size alone is an incomplete measure of recovery^12^. Genetic diversity is declining globally across many threatened and managed populations^13–15^, and future losses may continue even if habitat or population declines are successfully halted, because genomic erosion can lag behind demographic collapse^16,17^. Previously bottlenecked populations may therefore carry a drift debt, whereby genetic diversity continues to decline after population numbers recover, homozygosity increases, and harmful variants become increasingly exposed by inbreeding^18–20^. The key issue is whether populations that rebound numerically are also rescued from hidden genetic risk, or whether they remain on a delayed trajectory of genomic deterioration.

The severity and trajectory of the drift debt largely depend on demographic history. Long-term small populations may have already exposed some recessive deleterious alleles to selection, reducing the amount of standing deleterious variation and buffering subsequent inbreeding depression after bottlenecks ^21,20,22^. This might explain why some naturally rare or historically small populations appear to persist despite low diversity. The same process, however, can leave populations with long-term genetic risks from elevated relatedness, drift load, and limited adaptive potential^20,23^. The critical unresolved question is therefore whether long-term small populations that are rescued from extinction are genuinely buffered from genetic risk, or whether purging and continuing genomic erosion can unfold together after demographic recovery.

Resolving these alternatives is crucial to how we view species recovery, but remains elusive because it requires temporal genomic records that span collapse and recovery, and that link demographic history, conservation intervention, genomic change, and fitness consequences^24,25^. The Mauritius kestrel (*Falco punctatus*) offers a rare opportunity for addressing these questions. Endemic to Mauritius and once widespread on the island, the species experienced an extreme bottleneck, declining to only four known wild birds in 1974 following habitat loss and pesticide contamination, especially DDT^26,27^. Intensive conservation, including captive breeding, hand-rearing and release, supplementary feeding, nest-box provision and predator control, prevented extinction and enabled demographic recovery^28–30^. Alongside the remnant Black River Gorges population in the west, a second population was established in the Bambou Mountains (east Mauritius) in the 1980s. Subsequent monitoring showed that reintroduced populations followed contrasting trajectories under continued management^31^. The species therefore represents a rare case in which pre-collapse history, near-extinction, demographic recovery, and post-recovery dynamics can be reconstructed across time using time series genomic, ecological and management data. Because the Mauritius kestrel collapsed and was rescued decades ago, it also offers a rare window into the potential future for other rescued species now entering similar trajectories.

To resolve genomic change and its consequences across the history of near-extinction and recovery of the Mauritius kestrel, we sequenced 130 genomes spanning nearly 200 years, including pre-bottleneck historical museum specimens, individuals sampled through the early, middle and late post-bottleneck stages (Fig. 1d,e; Supplementary Table 1), and six other island and continental kestrel (*Falco*) species for comparative context (Supplementary Table 2). By combining temporal genomics with lifetime reproductive success data and forward simulations, we tested whether demographic rescue was accompanied by genomic recovery. We then asked whether long-term small effective population size buffered the species from inbreeding depression, or whether the recent bottleneck created a persistent genomic burden despite ongoing conservation management.

**Fig. 1.**
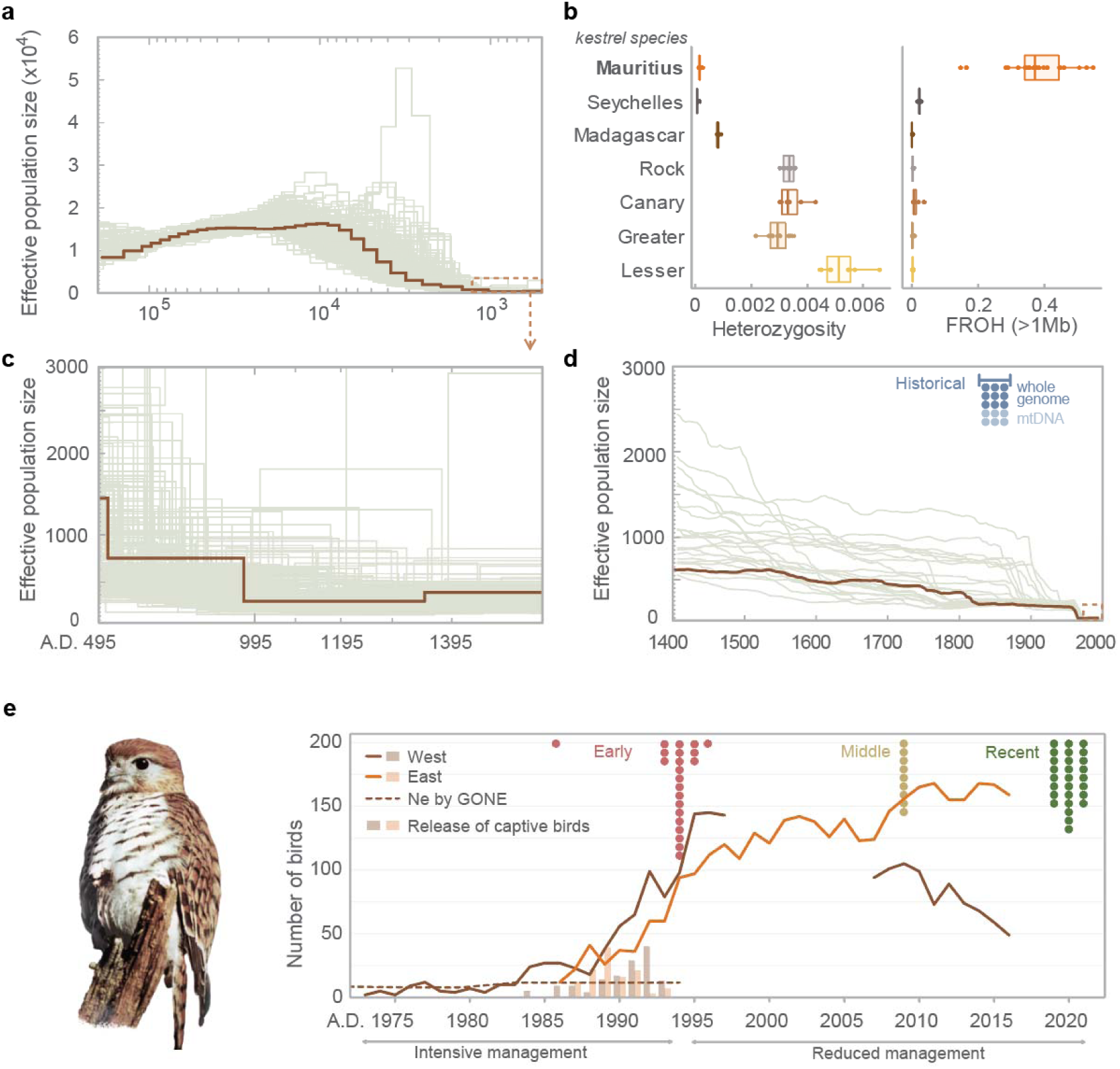
Demographic history and genetic status of the Mauritius kestrel. **a**, PSMC inference shows a long-term small effective population size. The dashed square in **a** shows the range in **b**. The brown line represents the highest-coverage genome; light green lines show bootstrap replicates from high-depth individuals. **b**, the small effective population size persisted prior to the human settlement of Mauritius. **c**, The Mauritius kestrel has low genetic diversity relative to other kestrel species, similar to the island species, Seychelles and Madagascar kestrels, but shows exceptionally high inbreeding. **d**, GONE2 inference indicates population decline over recent centuries, followed by a sharp mid-20th century crash. Brown line: inference using all early samples; light green lines: jackknife replicates. The Blue line marks the time period of the historical samples (dots). The dashed square represents the range of Ne shown in **e**. **e**, Recent population size and conservation history, with temporally sampled individuals indicated by dots. Photo of a captive female Mauritius kestrel “Queenie” released in 1992.

## Results

### Long-term small population size buffered, but did not remove genetic risk

The Mauritius kestrel has a long history of a small effective population size followed by a recent decline in the west population^32^. To characterise the demographic changes before the bottleneck, we performed PSMC^33^ and GONE^34^ analyses with 22 early Mauritius kestrel samples sequenced at high depth (Supplementary Table 2). The species sustained a small effective population size (Ne < 2,000) for more than two thousand years (Fig. 1a,b), far predating the recent human-driven decline in Mauritius. This prolonged persistence at a small population size is consistent with ecological estimates of the island’s carrying capacity. Monogamous breeding pairs of the Mauritius kestrel occupy territories of approximately ~1 km²^28^, suggesting that the ~1,865 km² Mauritius island could have supported ~3,000 breeding individuals. Its more recent demographic history reveals a further collapse, with Ne declining from ~450 in the 17th century to <100 by the 1960s (Fig. 1d), consistent with severe deforestation in Mauritius^35^ and pesticide use between 1940s to 1970s.

The Mauritius kestrel has exceptionally low genomic heterozygosity (1.22 × 10□□), similar to that reported for other severely bottlenecked species such as the Seychelles magpie robin (*Copsychus sechellarum*)^36^, Hawaiian crow (*Corvus hawaiiensis*)^37^, vaquita (*Phocoena sinus*) ^21^, and kākāpō (*Strigops habroptilus*)^38^, and only ~3% of that observed in the continental kestrel species studied (Fig. 1c). Its sister species, the Seychelles kestrel (*Falco araea*) and Madagascar kestrel (*Falco newtoni*), also show low genetic diversity, consistent with long-term demographic constraint in island lineages. In contrast, the Mauritius kestrel stands out for its pronounced inbreeding, with F_ROH_ inferred from runs of homozygosity longer than 1 Mb reaching as high as 0.39, far exceeding that of its sister island species (Fig. 1b). Together, these comparisons suggest that the exceptionally low diversity of the Mauritius kestrel largely reflects its long-term demographic history, whereas its extreme inbreeding reflects the recent severe bottleneck.

Comparative analyses indicated that insular kestrels carry fewer putative heterozygous deleterious alleles (Extended Data Fig. 1), used here as a proxy for masked load, the hidden component of genetic load not expressed in fitness effects^19^. Conversely, these insular kestrels carry more putative homozygous deleterious alleles, used here as a proxy for realised load, the expressed component of genetic load that reduces fitness (Extended Data Fig. 1). This pattern was consistent across two complementary annotations of deleterious variation, based on SnpEff^39^ loss-of-function (LOF) predictions and chicken CADD scores^40,41^ lifted to kestrel genomes; variants were polarised to the ancestral state across the kestrel phylogeny, and individual counts were normalised by derived neutral alleles. Together, these results suggest that the Mauritius kestrel had already exposed part of its deleterious variation before the recent bottleneck, potentially reducing the amount of masked to realised load conversion during subsequent severe inbreeding in the collapsed population^21,42^.

### Demographic recovery did not halt genomic erosion

To understand the temporal dynamics of genomic erosion through the population collapse and recovery, we sequenced one subfossil bone that likely dated to the 18th Century (Extended Data Fig. 2), 15 nineteenth-century museum specimens (“historical” samples, 1834–1875; nine whole genomes; Fig. 1d), and 59 post-bottleneck Mauritius kestrels sampled from 1986–1995, 2009, and 2019–2021 (Fig. 1e). Despite a 25-30 fold population decline, the reduction in genome-wide diversity across the bottleneck was modest relative to the severity of the demographic decline in the Mauritius kestrel. Heterozygosity decreased by only 18.4% between historical and early post-bottleneck samples (Fig. 2a), consistent with the already low baseline diversity in the 19th century. In contrast, mitochondrial diversity was completely lost, with only a single haplotype remaining (Extended Data Fig. 3). This contrast is expected because diversity metrics respond to bottlenecks at different rates: allelic diversity (including mtDNA haplotypes) is lost more rapidly than genome-wide heterozygosity^43^. Historical individuals already showed modest inbreeding (mean F_ROH_ = 0.07, ROHs > 1 Mb), together with an accumulation of short (100 Kb to 1 Mb) ROHs (Extended Data Fig. 4). After the bottleneck, F_ROH_ rose abruptly to 0.41 (Fig. 2b), accompanied by the appearance of long ROHs >10 Mb (Extended Data Fig. 5). Importantly, genomic erosion did not stop after demographic recovery, consistent with a continuing drift debt. Over the past ~30 years, corresponding to about seven generations, F_ROH_ increased further to 0.48, accompanied by an accelerated loss of genome-wide heterozygosity of 12.1%. Assuming this loss reflects genetic drift alone^44^, the decline corresponds to a harmonic mean Ne of ~27 over this interval.

**Fig. 2.**
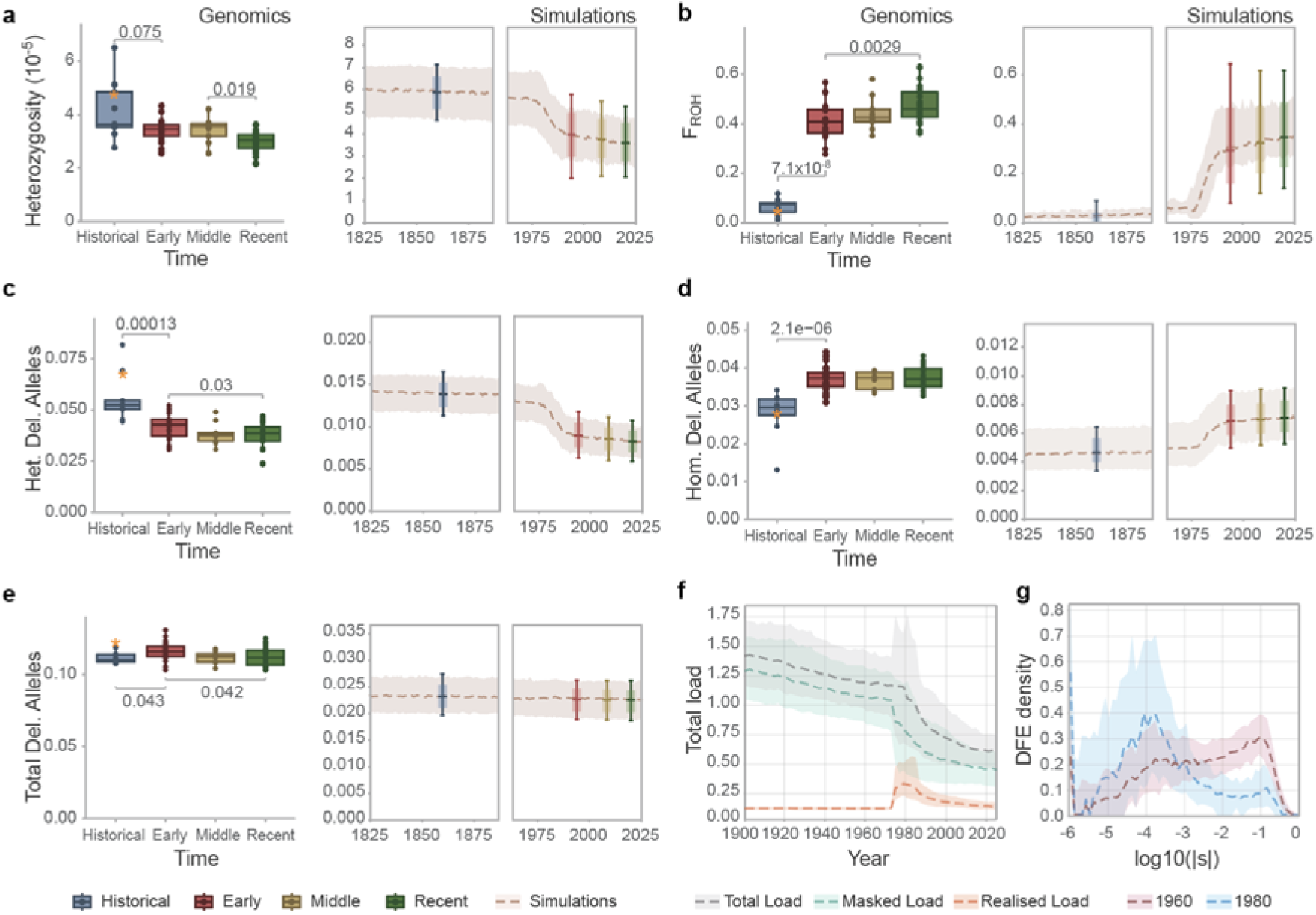
Temporal patterns of genomic erosion of the Mauritius kestrel. **a,** Continuous decline of genome-wide heterozygosity shown with both simulations and temporal genomic inferences. The orange star represents the subfossil bone sample. **b**, Increase in inbreeding through time. **C**, The corrected counts of heterozygous (Het. Del.) predicted by CADD score masked load in **f**. **d**, Corrected homozygous deleterious (Hom. Del.) corresponds with simulated realised load in **f**. **e**, Corrected counts of total deleterious alleles had different trend with simulated total load in **f**. **f**, simulated genetic load measured in predicted lethal equivalents. **g**, the shift of standing DFE before and after the bottleneck. The dashed lines show medians across replicate SLiM5 simulations and ribbons show the 10-90% range.

Although our temporal sampling spans approximately 200 years and captures the full course of the bottleneck and recovery, it comprises only four discrete time points rather than a continuous record. Therefore, to resolve the dynamics of genomic change through time, we developed forward-time simulation models closely parameterized to the Mauritius kestrel (Supplementary Methods), incorporating inferred demographic trajectories, life-history traits (e.g., mating and reproduction) and conservation interventions. Models were then validated against the temporal genomic data using genome-wide diversity, F_ROH_, and deleterious-variation summaries (Fig. 2a,b). The empirical changes in heterozygous and homozygous deleterious alleles matched the simulations, indicating a decline in masked load and an increase in realised load through the bottleneck (Fig. 2c-f; Extended Data Fig. 6). This conversion of masked into realised load reflects an increase in homozygosity due to both genetic drift and inbreeding^20^. After demographic recovery, simulations suggest that the realised load stabilized, whereas the total number of deleterious alleles increased even as total genetic load was predicted to decline (Fig. 2e,f; Extended Data Fig. 6). These apparently opposing trends reflect the fact that deleterious allele counts are unweighted, whereas genetic load is weighted by fitness effect. We explored weighting deleterious allele counts by class to calculate proxies for genetic load and observed signatures of purging of genetic load after demographic recovery (Extended Data Fig. 7; see Supplementary text). Thus, purging and genomic erosion occurred simultaneously: purging reduced the contribution of stronger-effect deleterious variants to total load, while genetic drift continued to reduce heterozygosity, increase homozygosity, and allow weaker-effect deleterious alleles to persist or rise in frequency. This left the standing variation enriched for mildly deleterious alleles (Fig. 2g; Extended Data Fig. 8). Thus, purging and genomic erosion are not alternative outcomes of conservation rescue, but can unfold together as populations recover demographically.

### Genomic erosion led to reduced reproductive performance

We quantified lifetime reproductive success for 18 individuals for which we also had genome data. We defined LRS as the total number of fledglings produced per individual across their lifetime. We then fitted negative binomial generalized linear models (GLM) relating LRS to heterozygosity, F_ROH_, and homozygous deleterious allele burden (both prediction methods), while controlling for the number of breeding years and sex. Across all three models, LRS was significantly lower in individuals with greater genomic erosion (Fig. 3a-c, Supplementary Table 3). Specifically, individuals with lower genome-wide heterozygosity, higher F_ROH_, and higher homozygous deleterious burden produced fewer fledglings over their lifetime. As expected, the number of breeding years was positively associated with LRS, but sex was not significant (*p* > 0.4). Together, these results provide rare individual-level evidence that genomic erosion has measurable consequences on reproductive success, indicating ongoing inbreeding depression despite the species’ long history of small effective population size. For rescued species, such links between genomic erosion and individual reproductive performance are critical in order to monitor recovery while accounting for genetic risk.

**Fig. 3.**
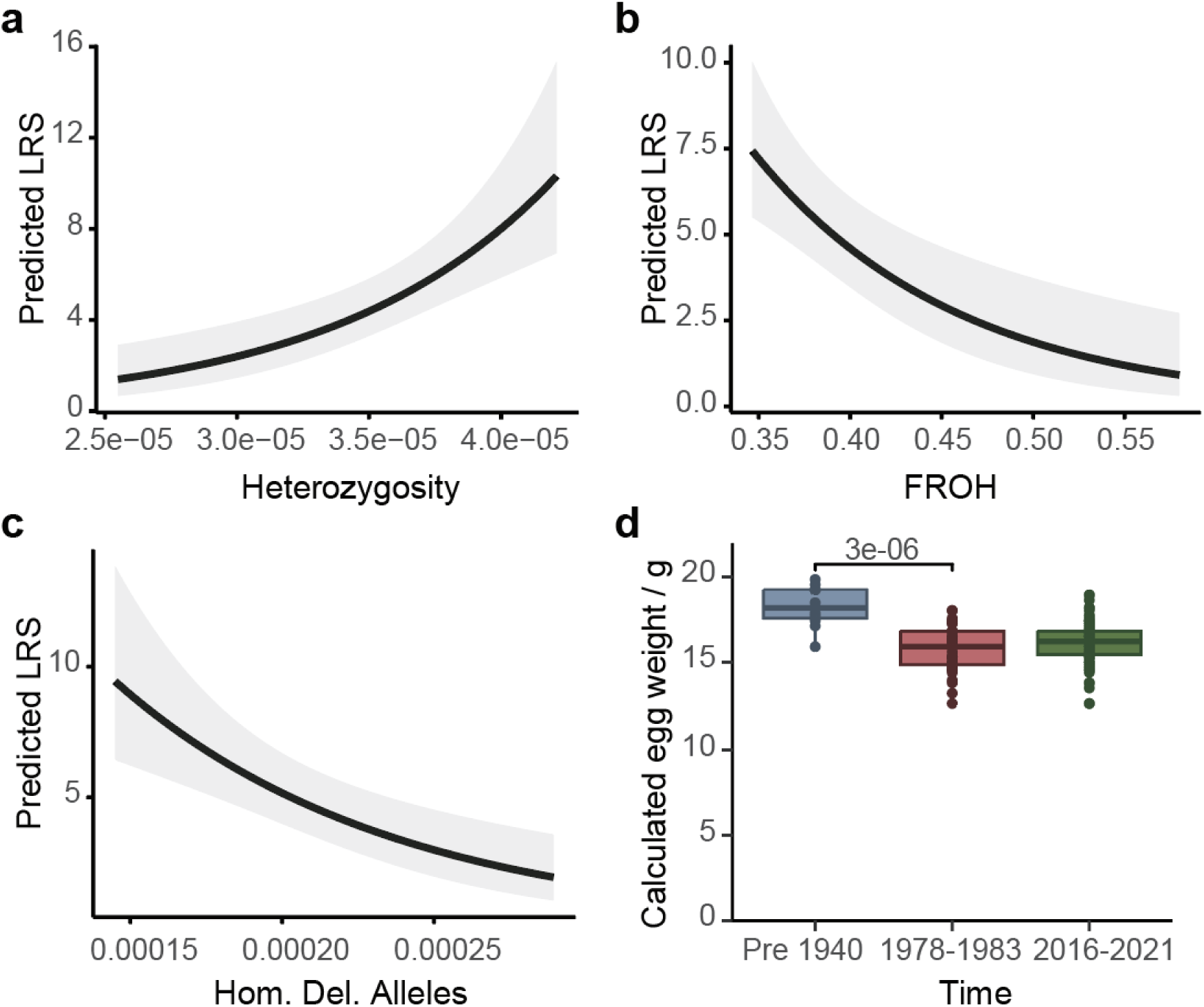
Reproductive performance, egg size, and genomic erosion in the Mauritius kestrel. **a**, Individuals with lower heterozygosity had lower predicted lifetime reproductive success (LRS) when controlling for the number of breeding years. **b**, Higher F_ROH_ was predicted to be associated with lower LRS. **c**, Higher number of homozygous deleterious alleles (LOF mutations) was predicted to be associated with lower LRS. All *p* < 0.001. Generalised linear model (GLM) was used as LRS data are typically zero-inflated. LRS was fitted as the response variable, using a negative binomial error distribution and a log link function. **d**, The egg weight declined for 17.8% through the bottleneck. The egg weight was calculated as *W = Kw(LB^2^)*. *Kw* = 0.0005248, a constant based on known Mauritius Kestrel egg fresh weights ^27^. *L* is egg length and *B* is egg breadth.

We observed a 17.8% decline of egg size through the bottleneck (Fig. 3d) using egg records from Jones 1987^27^ and newly collected data. Egg size is known to be positively correlated with hatching success in many birds including kestrel species^45,46^. Although DDT likely caused eggshell thinning in the Mauritius kestrel^26^, which is a well-established effect in birds^47,48^, observations and experiments showed that the egg size is not affected by DDT^47,49^. Consistent with this, egg size has not increased in recent years compared to values recorded in the 1970s–1980s (Fig. 3d), despite the reduction in pesticide use^26^. While ecological factors and founder effect cannot be ruled out, the persistent reduction of egg size provides independent evidence of reduced condition in the modern population, consistent with the genomic and LRS evidence for incomplete recovery of population health after the bottleneck.

### Conservation management sustains demographic but not genomic recovery

The Mauritius kestrel escaped extinction only through successful intensive conservation after the bottleneck^28,29^. Although current management is reduced, these interventions continue to have fitness benefits, as pairs breeding in nest boxes lay more eggs than those using natural nesting sites^32^. To evaluate how strongly the species’ persistence depended on conservation, we simulated two counterfactual scenarios under the same carrying capacity as the baseline model: one in which conservation management was never implemented, and another in which conservation management ceased in 2025 (Supplementary Table 4; see Supplementary Methods). These simulations show that historical conservation efforts were not only crucial, but remain necessary for persistence. In the absence of any conservation, all 100 simulation replicates went extinct within a few years (Fig. 4a). When management ceased in 2025, populations persisted longer but ultimately declined to extinction within approximately 35 years across all replicates (Fig. 4a).

**Fig. 4.**
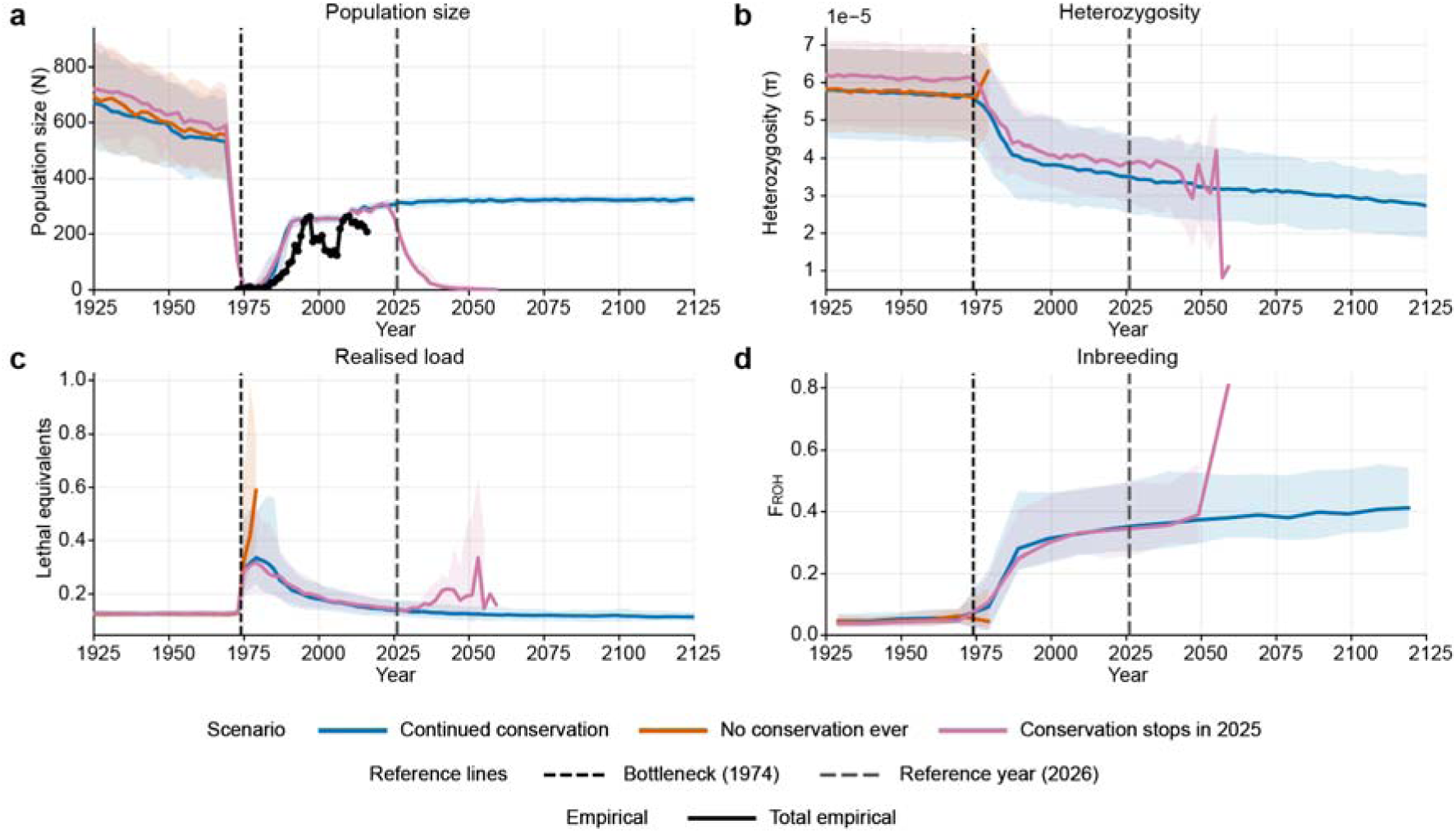
Simulated demographic and genomic trajectories of the Mauritius kestrel. **a**, Simulated trends of population size. **b**, Simulated trends of heterozygosity. **c**, Simulated trends of realised genetic load. **d**, Simulated trends of inbreeding measured as F_ROH_. All plotted through time under three management scenarios: continued conservation, no conservation ever and if conservation stopped in 2025. Lines show medians across replicate SLiM5 simulations and ribbons show the 10-90% range. The black line in **a** shows the combined empirical population trajectory. Vertical dashed lines indicate the year 1974, the year of the bottleneck, and 2026. The simulations recapitulate the post-bottleneck demographic trajectory and the accompanying genomic changes, including declining heterozygosity, increased realised load and F_ROH_. Continued conservation improves persistence relative to counterfactual scenarios, but drift debt creates conservation dependence after demographic rescue.

In effect, conservation management has likely buffered the fitness consequences of inbreeding depression by maintaining reproductive output at a sustainable level. If current conservation efforts are sustained, assuming a constant carrying capacity and stable environmental conditions, the short-term demographic outlook for the species appears stable (Fig. 4a). However, simulations predict a further decline in heterozygosity of approximately 23% over the next 100 years (Fig. 4b), consistent with a long-term drift debt^50^.

While realised load is projected to remain relatively stable (Fig. 4c), genomic recovery is not predicted. Heterozygosity continues to decline, and inbreeding is expected to gradually increase, with F_ROH_ staying above 0.4 on average. These projections indicate that continued management is critical to maintain population numbers, but it is unlikely to reverse genomic erosion or eliminate drift debt. This illustrates a central challenge for rescued species: management is important to maintain population numbers while the genomic consequences of past collapse continue to unfold.

## Discussion

Together, our results show that demographic rescue, purging, continuing genomic erosion, reduced reproductive performance, and conservation dependence can all unfold simultaneously in a rescued species. In the Mauritius kestrel, long-term small effective population size likely reduced masked load and softened immediate genetic consequences of collapse, consistent with expectations for populations that have persisted at small size^21,42^. Yet this demographic history did not prevent continued heterozygosity loss, rising homozygosity, fitness consequences consistent with ongoing inbreeding depression, or long-term dependence on management.^20,22,51^. Similar genetic buffering has been reported in other species with long-term small populations, including the vaquita (*Phocoena sinus*), Channel fox (*Urocyon littoralis*), and Alpine ibex (*Capra ibex*)^21,42,52^. However, the Mauritius kestrel shows that such buffering does not eliminate genetic risk. Rapid and intensive conservation management was nonetheless decisive in preventing extinction, rescuing the population from a demographic state in which stochastic processes alone could have driven loss when numbers were reduced to only a few individuals. Thus, long-term small effective population size may lessen some immediate genetic consequences of severe bottlenecks, but it does not remove the longer-term drift debt they create: demographic recovery was achieved, but genomic erosion continues, and long-term viability remains dependent on sustained management.

By integrating temporal genomics with long-term ecological data and empirically validated forward simulations, we show how demographic recovery and purging can coexist with continuing genomic erosion. Purging reduced total genetic load, but genetic drift continued to increase homozygosity, leaving a persistent drift debt. Although the short-term demographic outlook for the Mauritius kestrel appears stable under continued management, our results highlight that census trends alone do not reveal the ongoing loss of genetic diversity and rising homozygosity. Persistently low and declining genetic diversity may limit the species’ adaptive potential^53,54^, increasing vulnerability to future environmental changes^55^, such as introduced parasites, which are a threat to many other threatened species^35,56,57^. Moreover, sustained inbreeding may further reduce reproductive performance, potentially compromising long-term population viability^58,59^.

Our results suggest that conservation dependence can arise partly from genomic erosion, not only from ecological or demographic constraints^22,60^. Simulations show that the Mauritius kestrel remains conservation-dependent, predicting extinction if management ceases. This dependence is not explained only by current threats or limited habitat^28^, but is reinforced by genomic erosion and its association with reduced lifetime reproductive success (LRS). In other bottlenecked or rescued species, population recovery has also occurred without full genomic recovery, such as in the pink pigeon^12^ and whooping crane^61^. By contrast, some rescued populations may recover genetic resilience when demographic expansion is rapid and substantial^62^, indicating that the genomic outcome of demographic recovery is species specific. Indeed, studies on vaquita^21^, kākāpō^38^, and Iberian lynx^63^ show that the balance between genetic drift, purging, and realised load varies with demographic history. Conservation assessments that respond mainly to improved census size may therefore understate risk in some rescued species. This may already be true for the Mauritius kestrel, which has been downlisted on the IUCN Red List after demographic recovery and assessed as having low conservation dependence on the Green Status despite marked genetic depletion and continuing genomic erosion. For such species, downlisting may obscure continuing conservation needs while drift debt is still unfolding^22^. Thus, some rescued species may remain conservation-dependent because their past decline created a drift debt that drives continued genomic erosion.

These findings underscore the importance of preserving, and where possible restoring, genetic diversity in bottlenecked species^15,17^, and at the same time maintaining conservation actions that buffer populations whilst genomic risks continue to unfold. More interventionist approaches, such as interspecific hybridization or genome editing, remain subject to ongoing debate^64–67^. Yet these options may become increasingly relevant when traditional management can only maintain population size and cannot restore the genetic variation lost from the gene pool. For the Mauritius kestrel, such approaches remain prospective rather than immediately actionable. A more immediate option may be to establish a new captive population, where reproductive output could be increased under managed conditions and remaining unique genetic diversity preserved as both a demographic and genomic safeguard^61,68^. The persistence of drift debt highlights why genomic restoration may need to be considered alongside habitat restoration, nest-site management, and captive breeding. For populations rescued from the brink of extinction, lasting conservation success must therefore mean not only demographic recovery, but also accounting for genetic diversity loss and rising genetic load over the long term.

## Acknowledgments

We thank the curators and collection staff of the natural history museums that generously provided access to historical specimens and assisted with sampling for this study: the Natural History Museum, London (NHMUK); Muséum national d’Histoire naturelle, Paris (MNHN); the University Museum of Zoology, Cambridge (UMZC); the Museum of Comparative Zoology, Harvard University (MCZ); and Naturalis Biodiversity Center, Leiden. Samples of kestrel species were kindly provided by J. Carrillo (Canary Island kestrel), A. van Zyl (Rock, greater, lesser kestrel), and Réné De Roland Lily Arison/S. Goodman (Madagascar kestrel). Bioinformatics analyses were carried out at the Globe Institute Mjolnir High Performance Computer with support from Bent Petersen. We acknowledge the long-term monitoring and conservation efforts by the Mauritian Wildlife Foundation, and by the National Parks and Conservation Service of the Government of Mauritius, to recover Mauritius’ threatened birds.

This work was supported by the European Research Council (101078303) and the Swedish Research Council for Sustainable Development (2022-00536). Further support was obtained from a Danish National Research Foundation grant (DNRF143), a Royal Society International Collaboration Award (ICA/R1/201194); a Natural Environment Research Council Grant (NE/S007229/1). The Mauritian Wildlife Foundation is supported by Standard Bank (Mauritius). JG was supported by Research England’s Expanding Excellence in England (E3) Fund, UK Research and Innovation. AS was supported by a Leverhulme ‘Space for Nature’ Doctoral Scholarship.

## Author contributions

This project was led by H.M and X.W. with inputs from C.J., K.N., J.G., and C.vO.. H.M designed the study. A.S. (Anna Stuart) performed SLiM simulations. K.N. performed analyses related to LRS. S.H. and C.J. collected blood samples and ecological data. K.N., S.H., S.B., M.T.P.G., V.T., K.R., J.G. and H.M. did sample curation. S.D.N. performed DNA extraction and library preparation. C.P. performed mapping and quality check for historical samples. J.P.H. collected the subfossil bone sample. A.S. (Alex Strang), and M.W. performed DNA extraction and library preparation of the subfossil bone sample. C.vO. performed weighted load partitioning. X.W. performed all other analyses. X.W., A.S. (Anna Stuart), K.N., C.vO. and H.M. wrote the original draft and all authors edited and approved.

## Data and Code availability

Raw sequence reads have been deposited in the Sequence Read Archive (SRA) under BioProject PRJxxxxx. Scripts used for sequence data processing and analysis (https://github.com/XJWang42/mk) and scripts for simulations (https://github.com/as2376/Kestrel_SLiM) are available on Github.

## Methods

### Samples and sequencing

For modern samples, blood or tissue samples were taken and preserved in 95% Ethanol. Samples were extracted using a DNeasy Blood and Tissue Kit (Qiagen, Hilden, Germany) and submitted for 150bp paired-end sequencing in an S4 Illumina NovaSeq 6000 at Novogene (Cambridge, UK). For the Mauritius kestrel (Table S1), 22 individuals from the ‘early’ time period (1986 - 1996), 13 from the west population and 9 from the east population, were sequenced at a high depth, 45.6x on average (39.1x - 65.8x). The remaining 37 samples, all sampled from the east population, including 1 early-, 9 middle- (2010) and 27 recent-time period (2019-2021) samples, were sequenced at a lower depth, 8.6x on average (7.9x - 10.2x). The other kestrel species were sequenced at an average depth of 21.9x, ranging from 17.2x to 33.2x (Supplementary Table 2). Only the high-depth Mauritius kestrel samples were used for interspecies comparisons.

For historical samples, 15 individuals were processed from toe-pad tissue obtained from Natural History collections and one subfossil bone sample (Supplementary Table 1). The museum specimens were from the 19th Century with uncertainties of the sampling year of four samples, but presumably all in the range of 1832 to 1875 (Supplementary Table 1). Historical toe-pad samples were extracted for ancient DNA following Gilbert et al.^69^ in ultra-clean laboratories at the University of Copenhagen. Three genomic libraries per-extraction were built following Kapp et al.^70^ and amplified as informed by qPCR results. All samples were submitted for 150bp paired-end sequencing in an S4 Illumina NovaSeq 6000 at Novogene (Cambridge, UK).

The subfossil bone sample was collected from a cave site excavation near La Prairie by Julian P. Hume, Mauritius (2006-2009) on private land owned by Owen Griffiths and housed at NHM (Tring, UK). It was a tibia tarsus from an unarticulated specimen, identified as part of a Mauritius kestrel. The bone was grinded into powder and bleached. 500ul of bleach (0.5%) was added to the bone powder and incubated at 23 °C with rotation for 10 mins. The sample was centrifuged for 2 mins, supernatant discarded. The pellet was washed 3 times with 500ul DD water, with vortexing and 2-minute centrifugation between each wash. DNA extraction was done following Dabney et al 2013, modified by replacing the Zymo Spin V column with Roche High Pure Vital Nucleic Acid Spin columns and two elution steps of 50 ul TET buffer. Three DS-stranded libraries were prepared following the building protocol outlined by Meyer and Kircher ^71^ with dual indexes. Sample Screening was done by the NHM on both the AVITI and NovaSeq. The libraries were sequenced as 150-bp paired-end reads with Illumina NovaSeq 6000 at Novogene.

### Modern sample mapping

For all samples, reads were mapped to the reference genome of the Mauritius kestrel (GCF_963210335.1). The reads of the modern samples were mapped using GPU-based BWA-MEM^72^ implemented in Parabricks Accelerated Genomics Pipeline 4.0.0^73^ with default parameters. To match with the low-depth samples in temporal analyses of the Mauritius kestrels, samples with high depths were downsampled to ~9x by randomly keeping 20% of the reads in the BAM file with Samtools 1.10^74^.

### Historical sample mapping

For historical samples and the subfossil osteological sample, raw reads were first filtered and merged with SeqPrep2 (https://github.com/jeizenga/SeqPrep2). The first and last five base pairs of the reads of the subfossil sample were masked. The merged longer reads were mapped to the reference genome using BWA-ALN 0.7.17^75^ with the following parameters: “-l 1024 -n 0.03 -o 2”. The DNA damage patterns were examined with mapDamage 2.2.2.

### Variant calling

Adapted from GATK Best Practice Workflows^76^, first, a raw GVCF for each sample was generated using GPU-based GATK HaplotypeCaller implemented in Parabricks Accelerated Genomics Pipeline. Second, GVCF files were merged into genome databases using GATK (4.4.0.0) GenomicsDBImport. Last, joint genotyping was performed with GATK GenotypeGVCFs.

### Variant filtering

The following filters were applied to all variants for all downstream analyses: “QD<2.0, SOR>3.0, FS>60.0, MQ<40.0, MQRankSum<-12.5, ReadPosRankSum<-8.0, AC>1”. Only SNPs on autosomes and scaffolds longer than 500kb were retained. The variants of the low-depth samples were filtered with a minimum GQ of 12, a minimum depth of 4 and a maximum depth of 30. The variants of the high-depth samples were filtered with a minimum GQ of 20, a minimum depth of 6 and a maximum depth of 120. Only biallelic SNPs were retained for further analyses. For temporal analyses including historical samples with relatively higher mismatch rates (Supplementary Fig. 1), transitions were removed to obtain high-quality SNPs.

All samples were additionally mapped to the reference genome of the lesser kestrel (*Falco naumanni*, GCF_017639655.2), and variant calling and filtering were performed using the same pipeline as described above. We used an additional reference genome for two purposes. First, we observed significantly higher numbers (p<0.001) of autosomal heterozygous sites in female Mauritius kestrels when mapping to the male Mauritius kestrel reference, which has no W chromosome (Supplementary Fig. 2a), but no such bias (p=0.34) when mapped to the female lesser kestrel reference (Supplementary Fig. 2b), with no mapping quality observed between the sexes (Supplementary Fig. 2c,d). When removing the reads mapped to the W chromosome of the lesser kestrel from the assemblies mapped to the Mauritius kestrel, this bias was controlled (p=0.063, Supplementary Fig. 2e). Thus, for all downstream analyses, we masked the SNPs covered by the reads mapped to the W chromosome of the lesser kestrel. Second, the variants called with the reference genome of the lesser kestrel were specifically used for the annotation of functional variants using SnpEff (see below). The gene annotation of the Mauritius kestrel genome contains approximately 20% fewer annotated genes compared to that of the lesser kestrel, which may reduce the reliability and completeness of functional variant annotation, thus we used the genome of the lesser kestrel instead for functional annotations by SnpEff.

### Mitochondrial sequence reconstruction

To reconstruct the mitochondrial DNA sequences, reads that mapped to the mitochondrial reference assembly were extracted using Samtools 1.10. Variant calling was performed using BCFTools mpileup (version 1.21, REF) and BCFTools call under a haploid model. Called variants filtered with a quality score (QUAL) > 50 and a mapping quality (MQ) > 40. The filtered variants were further manually checked with IGV 2.19.3^77^ in the BAM files. The haplotype network was generated with PopArt 1.7 ^78^ using Minimum Spanning methods.

### Heterozygosity and runs of homozygosity

The number of heterozygous sites were counted for each sample with BCFTools, and genome-wide heterozygosity was calculated by dividing the number of heterozygous sites with the number of callable sites, counted from the GVCF file. The runs of homozygosity were identified with BCFTools with default parameters. We calculated F_ROH_ as the inbreeding rate, by dividing the total length of ROHs longer than 1 Mb by the total length of the reference genome.

### Potential population difference

To test for potential population structure, we performed PCA analysis with PLINK v1.9.0-b.8^79^ for the early samples, using SNPs with a missing rate lower than 10% from the high-depth dataset (22 early Mauritius kestrels and other kestrel species). The two populations of Mauritius kestrel, east and west, had no population structure, and no difference in genomic heterozygosity and FROH (Supplementary Fig. 3). This is expected as the east population is established by introducing individuals from the west population from 1983 to 1993, overlapping with the sampling time. Thus, we didn’t differentiate the two populations further in genomic analyses.

### Long demographic history

Long-term demographic history (500 - 200,000 years ago) was inferred using the Pairwise Sequentially Markovian Coalescent (PSMC) model^33^ with an individual sampled in 1995 at high depth (65.8x). Consensus sequences were generated from mapped reads using Samtools, applying minimum mapping and base quality thresholds of 30 and a minimum depth of 5 (--min-MQ 30, --min-BQ 30, -d 5). The outputs were converted into PSMC input format using fq2psmcfa with a quality filter of 30. To further determine the robustness of PSMC results, we performed 10 rounds of bootstrapping on each of the high-depth Mauritius kestrel samples following PSMC’s recommendation. Analyses were performed with parameters -N30 -t5 -r5 -p “1+1+1+1+30*2+4+6+10”^80^. We used a generation time of four years and a mutation rate of 6.29×10^−9^ per generation, converted from the per-year mutation rate of peregrine and saker falcons^81^.

### Recent demographic history

Recent demographic history (since 1400) was inferred using GONE2^34^, with a recombination rate parameter of 2 (-r 2) and disabling the genetic map flag (-g 0). To obtain confidence intervals, a jackknife approach was implemented by iteratively removing one individual at a time from the dataset.

### Annotation of putatively deleterious variants

We used two methods to annotate putatively deleterious mutations. First, we used SnpEff 5.2^39^ to annotate loss-of-function (LOF), missense mutations, and mutations outside of exons based on the reference genome and gene annotation of the lesser kestrel (GCF_017639655.2). Second, we assessed the deleteriousness of the mutations using Combined Annotation Dependent Depletion (CADD) scores using LoadLift pipeline^68^. We did liftover of the chicken (*Gallus gallus*) CADD scores^41^ based on reference genome Galgal6 to the reference genome of the Mauritius kestrel.

In detail, to enable coordinate conversion between the reference genome of chicken and the Mauritius kestrel, whole-genome pairwise alignments were first constructed and converted into chain files. Both genomes were converted into UCSC two-bit format using *faToTwoBit* from the UCSC source code^82^ and were split into smaller units to facilitate parallel alignment. The chicken genome was split by chromosome, while the genome of the Mauritius kestrel was partitioned into sequence chunks using *faSplit*. Pairwise alignments between all combinations were computed using lastz^83^ with parameters optimized for sensitive long-range alignments (--hspthresh=2200 --inner=2000 --ydrop=3400 --gappedthresh=10000). Alignments were converted into chain format using *axtChain*, applying a minimum alignment score of 5000 and a loose linear gap model. All chain files were merged with *chainMergeSort* and sorted with *chainSort* to produce a genome-wide chain set. To refine alignments, chain files were netted using *chainNet*, and a subset of high-confidence chains was extracted using *netChainSubset*. Chicken CADD scores were lifted to the Mauritius kestrel genome using CrossMap^84^ with the generated chain files.

### Inference of ancestral states

To infer the ancestral state of the variants, we used IQ-TREE 2.4.0^85^ to reconstruct a maximum likelihood (ML) phylogenetic tree using the SNP alignments of high-depth kestrel genomes. An initial model-selection step identified the best-fitting substitution model, and TVM+F+G4 was selected for downstream analyses. For each chromosome, ML trees were generated using ultrafast bootstrap approximation with 1,000 replicates, and combined into a genome-wide phylogenetic tree. Branch lengths were then added to the topological tree using all chromosomal alignments to produce the final consensus tree.

The consensus tree was used as the guide to infer the ancestral states of the loci from the ancestral node of all kestrel samples. For each site, the maximum posterior probability across all possible nucleotide states was calculated. 97.0% Sites were retained with the maximum probability ≥0.95, indicating high confidence in the inferred ancestral state.

### Inference and normalization of genetic load based on genomic data

We estimated genetic load based on the annotations of putatively deleterious alleles with both SnpEff and CADD methods. For functional annotation with SnpEff, LOF and missense variants in which the reference allele matched the ancestral state were retained as putatively deleterious variants. Intergenic variants and intronic variants for which the reference state matched the ancestral state were used as neutral variants. For CADD annotation, the top 5% most deleterious predicted variants (CADD>13.0) were used as putatively deleterious variants, and the 50% less deleterious variants (CADD<3.0) were used as neutral variants. Because CADD estimates deleteriousness for all three possible mutations, SNPs with either the reference or alternative allele matching the ancestral state, and with liftover CADD scores available, were retained. The CADD score for the corresponding derived mutation was then used as the variant’s CADD score. Fixed variants within the Mauritius kestrel were removed. To account for differences in SNP quality among samples, raw counts of derived deleterious alleles were normalized by the total number of derived neutral alleles in each sample. Normalized counts of homozygous deleterious alleles were used as an approximation of genomic realized load, normalized counts of heterozygous deleterious alleles as an approximation of masked load, and normalized counts of total deleterious alleles as an approximation of total load.

### Lifetime Reproductive Success

To investigate the potential ongoing fitness consequences of genome erosion, we compiled lifetime reproductive success (LRS) data for 18 individual Mauritius kestrels for which we also had genome data, using the long-term breeding and resightings data from 1980s onwards ^32^. Briefly, during each breeding season, nesting attempts were monitored from egg laying to fledging, adult birds were identified by their unique colour rings, and chicks were colour-ringed before leaving the nest to enable subsequent identification in the field. Using these data, we estimated LRS as the total number of fledglings produced over an individual’s lifetime for a sample of Mauritius kestrels. Of these, three individuals were known to be alive in the 2024/25 breeding season, so their LRS may have been under-estimated. As a result, we also generated a weighting variable for our models in which an individual was included as 1 (dead) if not resighted in the 2024/25 breeding season and 0 (alive) if it was resighted.

### GLM with genetic estimates

To quantify the relationships between LRS and our measures of genome erosion, we used generalised linear models (GLMs) in the statistical programming language R^86^. Since LRS data are typically zero-inflated, we fitted GLMs with LRS as the response variable, assuming a negative binomial error distribution, and using a log link function. We set-up separate models each with a different measure of genome erosion as a predictor variable – genome heterozygosity (het), FROH and corrected derived homozygous deleterious alleles (LOF or top 5% CADD). LRS in wild birds often correlates with longevity^87^, so we also included the number of breeding years for each individual as a predictor variable in each model. Lastly, we also included individual sex as a predictor to check for any sex-specific effects on LRS. For each measure of genome erosion, we ran two GLMs; one including all individuals and another limited only to those individuals known to have died by the 2024/25 breeding season.

Initial results suggested rescaling the measures of genomic erosion would be beneficial, so we rescaled these in the GLMs using the “*scale*” function. With this adjustment, all models converged properly with no warnings. The results were identical whether or not we weighted out individuals known to be alive in the 2024/25 breeding season, so for brevity we simply report the results of the weighted models.

### SLiM simulations

Forward-in-time individual-based simulations were used to model demographic and genomic change in the Mauritius Kestrel through historical decline. Simulations were implemented in SLiM v5.1 using the non-Wright-Fisher (non-WF) framework^88,89^. This allowed an annual, age-structured life cycle with overlapping generations, sex-specific monogamous pairing, stochastic fecundity, density-dependent survival and managed releases^21,50^. The model was designed as an empirically anchored demo-genetic process model. Its purpose was to propagate the known demographic history, life-history structure and conservation interventions and then compare simulated genomic trajectories with empirical temporal genomic data. See Supplementary Methods for details.

---

**Extended Data Fig. 1.**
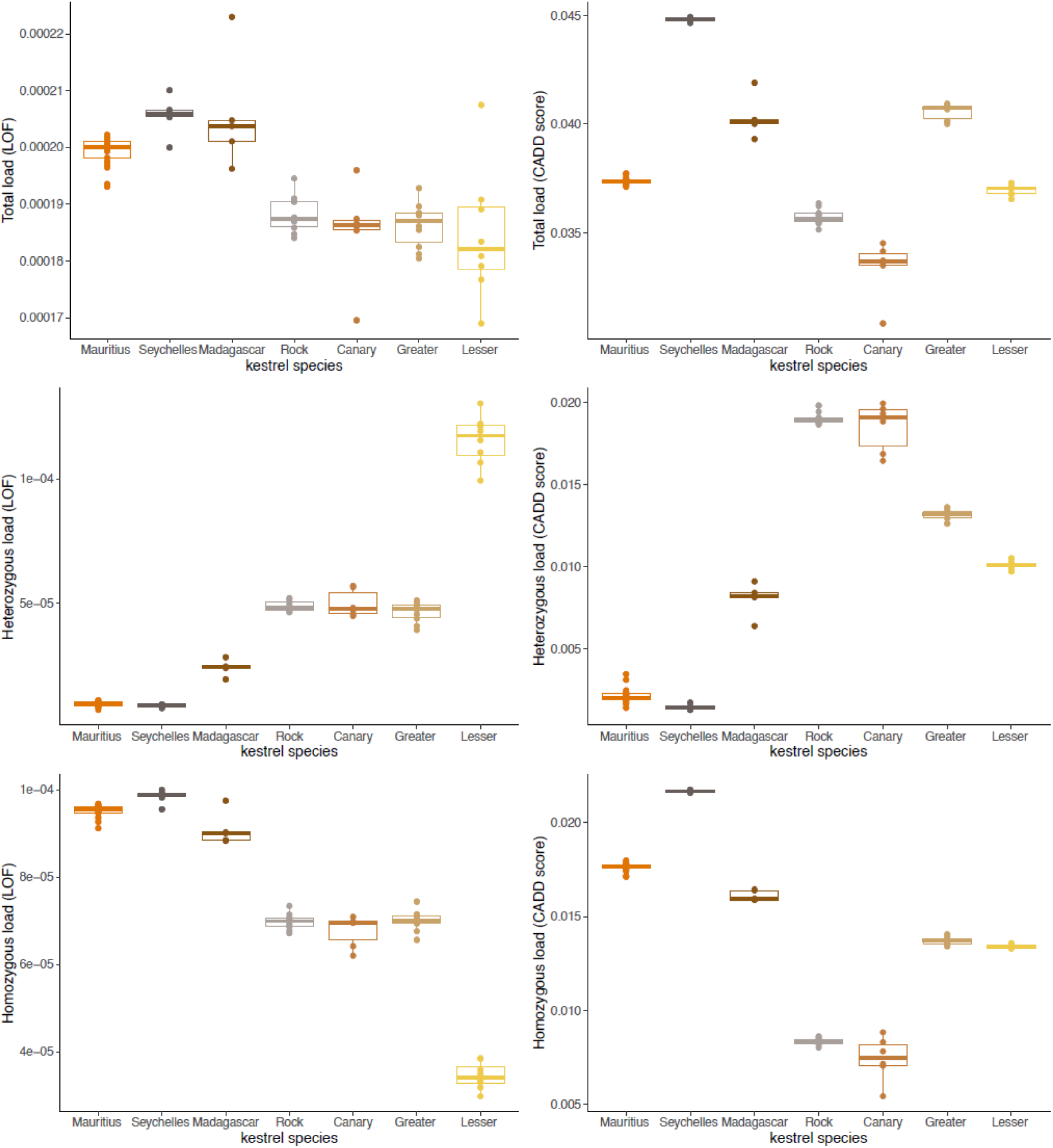
Interspecific comparisons of genetic load. The estimates were based on two methods of annotation of deleterious variation: SnpEff loss-of-function (LOF) predictions and chicken CADD scores lifted to the lesser kestrel genome. Raw counts of derived sites were corrected by the number of neutral derived alleles. Island species carry fewer putative heterozygous deleterious alleles and more homozygous deleterious alleles.

**Extended Data Fig. 2.**
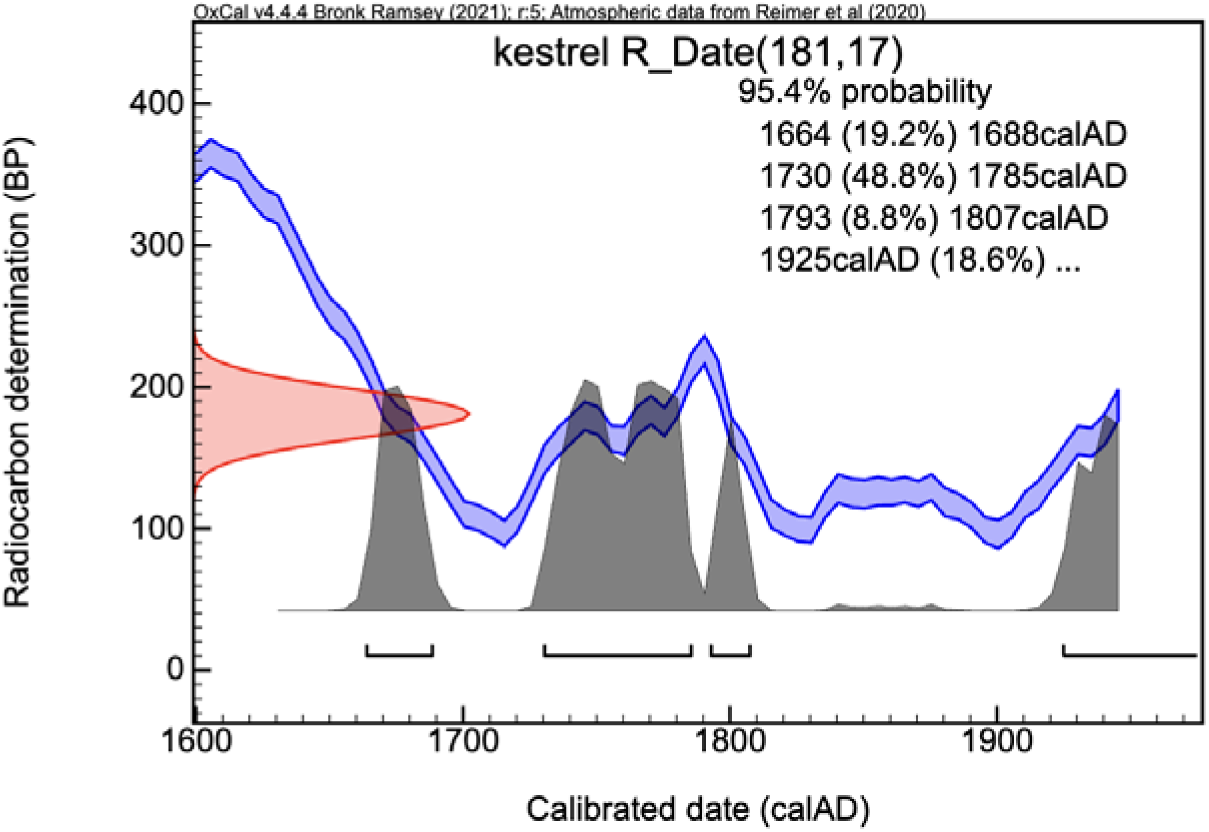
Dating of the subfossil bone. The radiocarbon date of the sequenced subfossil bone was 181±17 years ago from 1950 AD, with a possibility of 68% earlier than 1785 AD.

**Extended Data Fig. 3.**
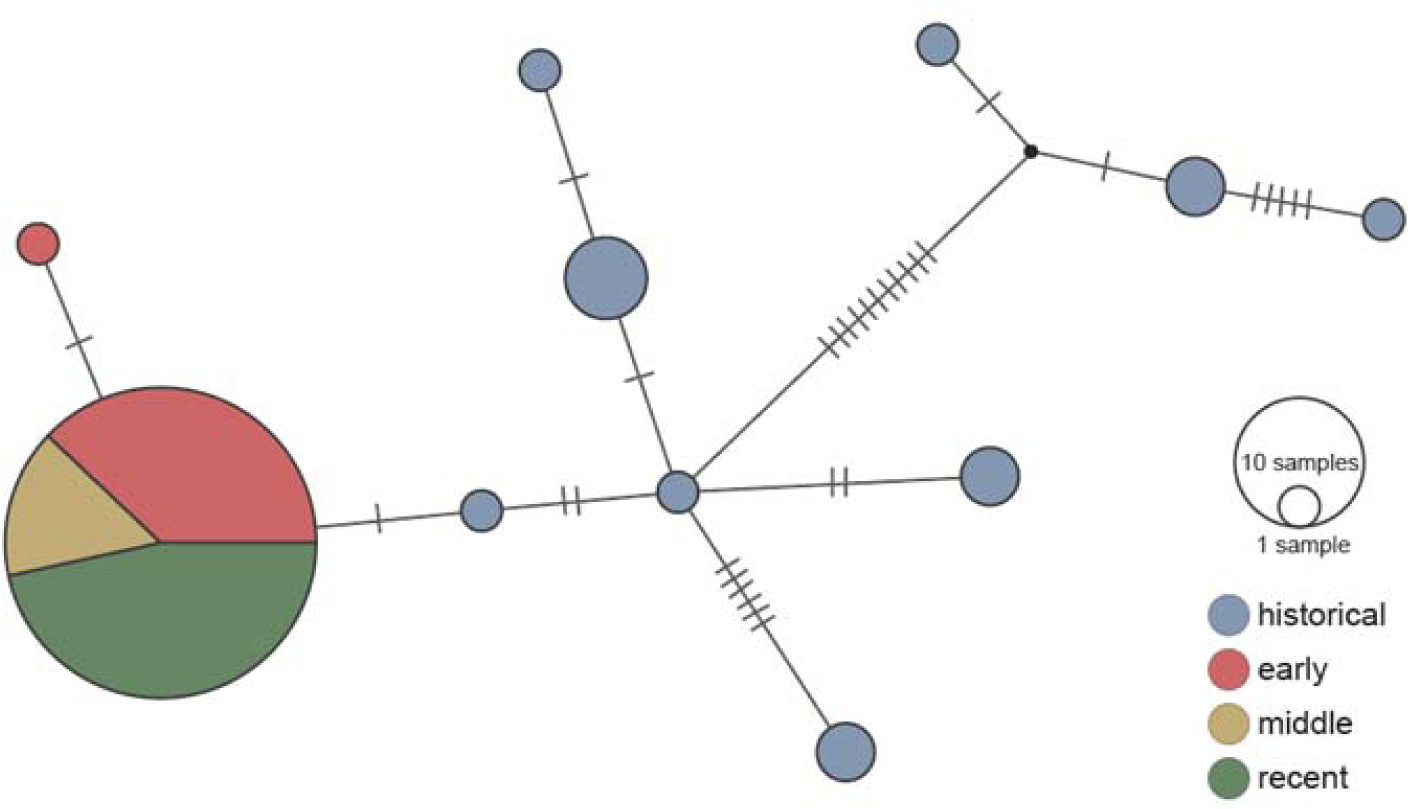
Haplotype network of mtDNA sequences of the Mauritius kestrel. Each circle represents an individual haplotype. Short lines represent substitutions between haplotypes. The small red circle on the left represents a haplotype from a single male sampled in 1994.

**Extended Data Fig. 4.**
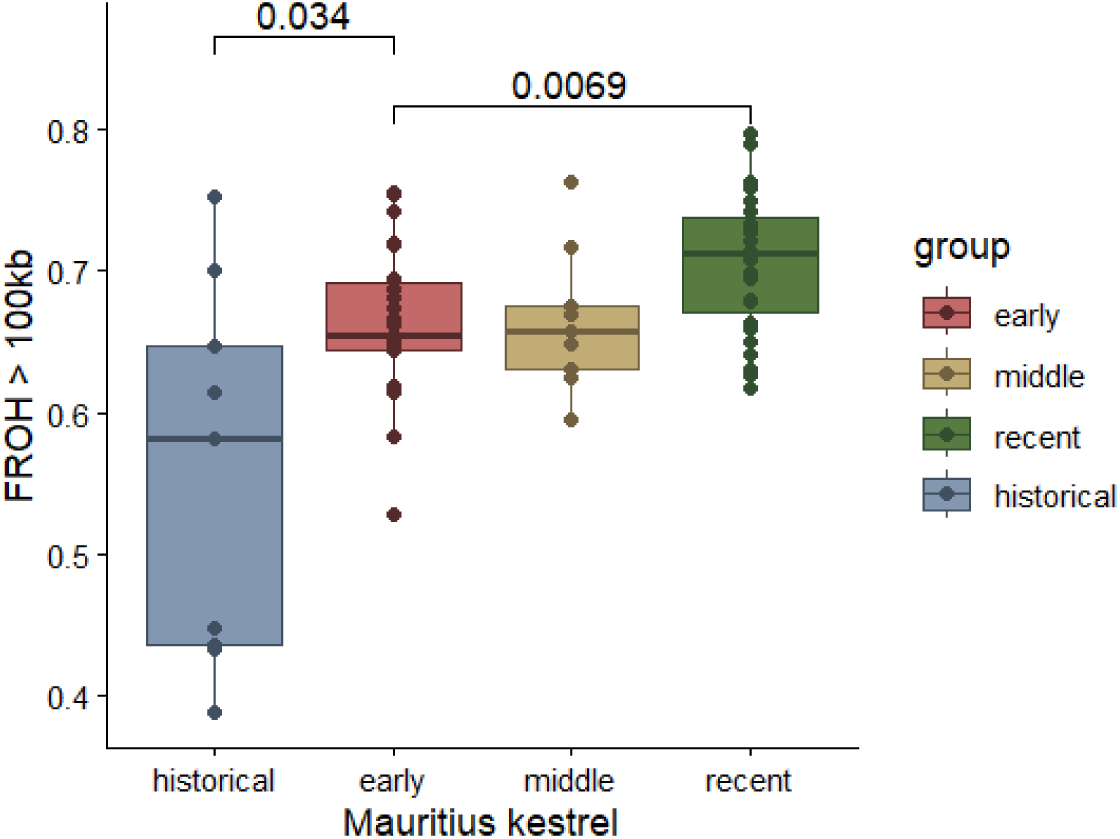
Temporal comparisons of inbreeding in the Mauritius kestrel. F_ROH_ with ROHs > 100 kb showing the historical F_ROH_ including short ROHs before the bottleneck was already above 0.5 on average.

**Extended Data Fig. 5.**
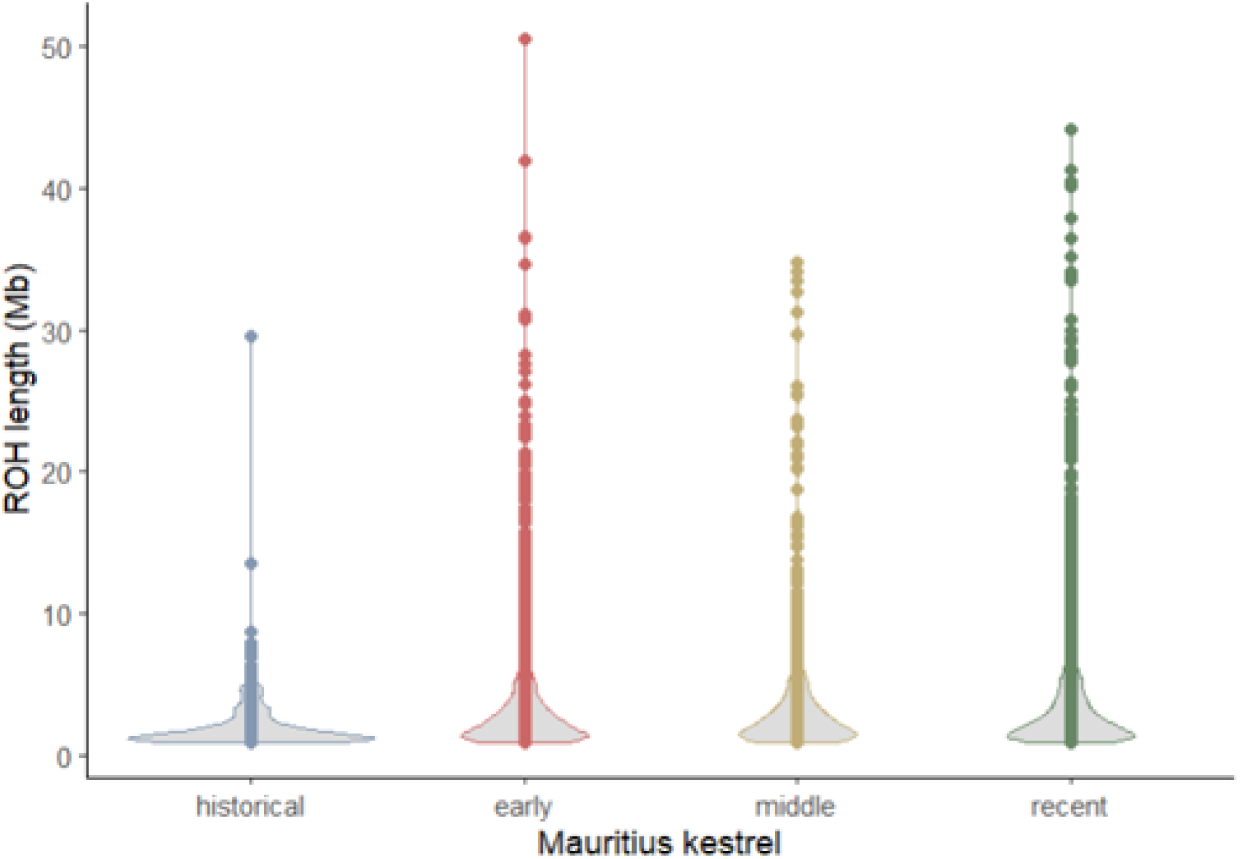
Distribution of lengths of ROHs in the Mauritius kestrel. Each dot represents an ROH > 1 Mb. Post-bottleneck individuals accumulate more long ROHs >10 Mb.

**Extended Data Fig. 6.**
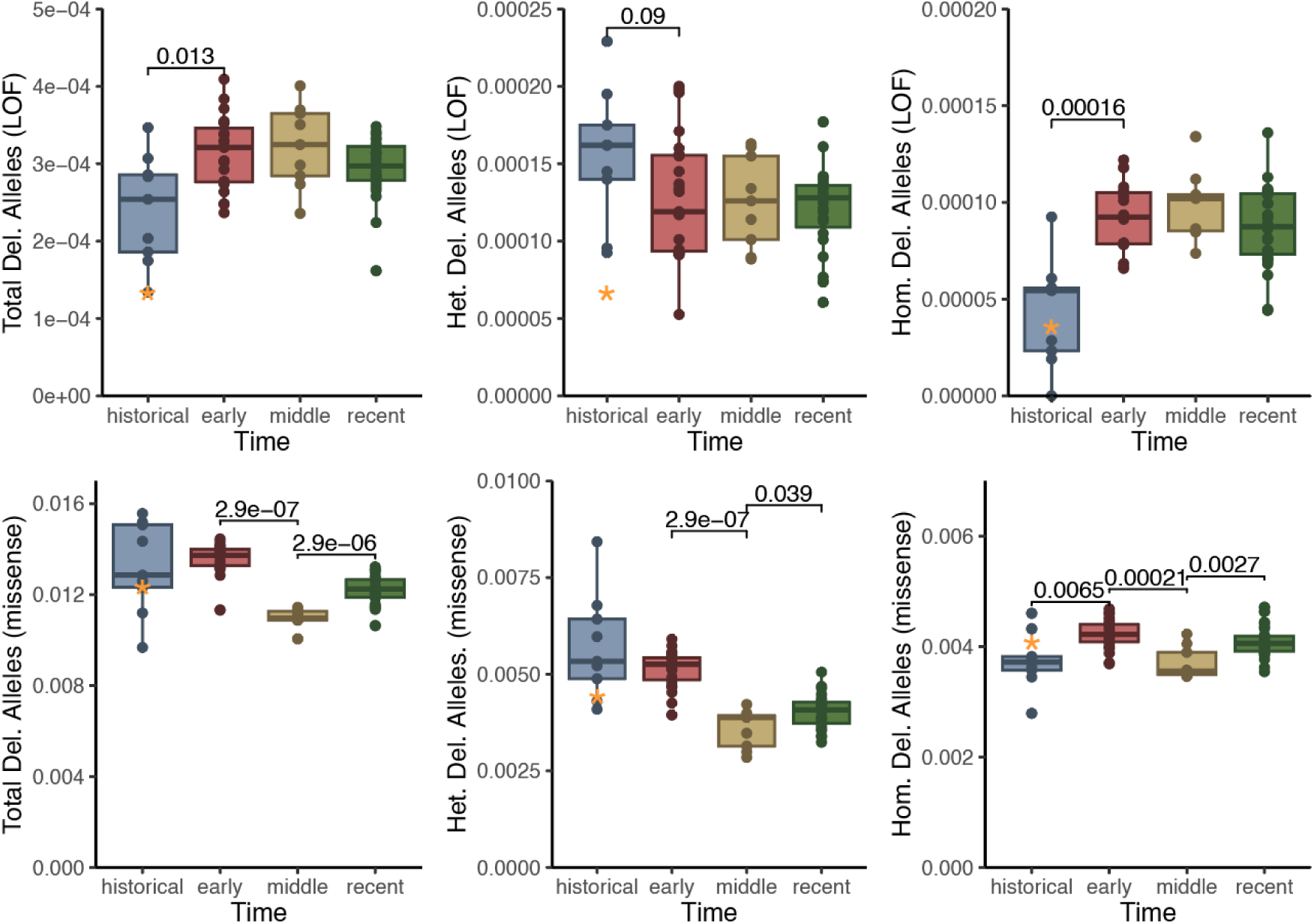
Temporal comparisons of estimates of genetic load based on SnpEff functional annotations. The derived Loss-of-Function (LOF) and missense mutations were considered as deleterious mutations. Raw counts of heterozygous sites (Het. Del.), homozygous sites (Hom. Del.) and total alleles (Total Del.) were corrected by the total number of derived neutral alleles for each individual. The orange star represents the subfossil sample.

**Extended Data Fig. 7.**
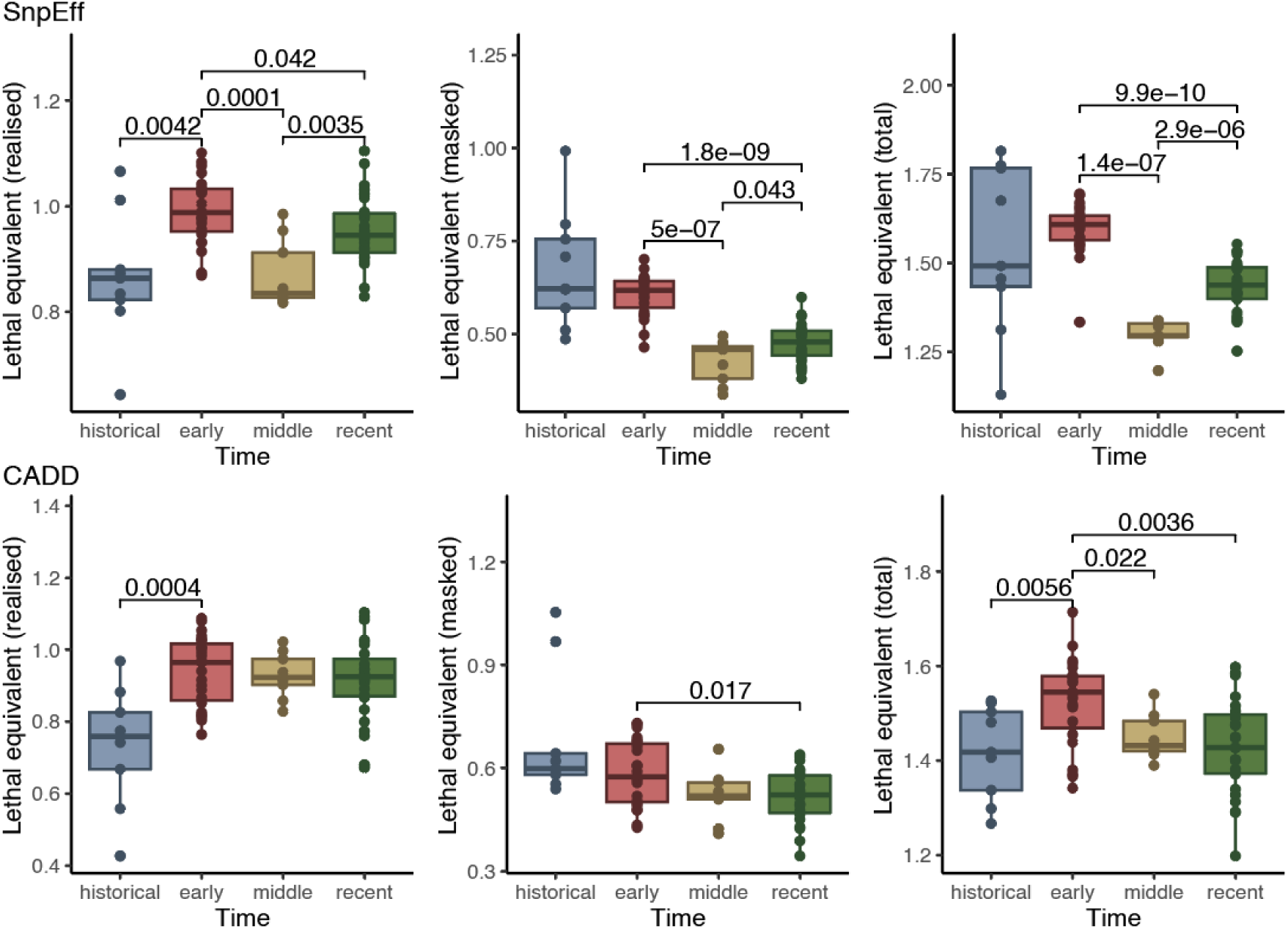
Genetic load estimates in approximate lethal-equivalent units using simulation-anchored class-specific coefficients. Realised load was higher in the early post-bottleneck group than in the historical group, consistent with conversion of previously masked recessive burden into homozygous realised burden. Masked load declined after the early post-bottleneck period, consistent with loss of heterozygosity and exposure of recessive variants. Total genetic load was highest in the early post-bottleneck group and lower in the middle group, consistent with partial purging of exposed deleterious burden. Total load was slightly higher again in recent samples than in the middle group, suggesting possible renewed drift load.

**Extended Data Fig. 8.**
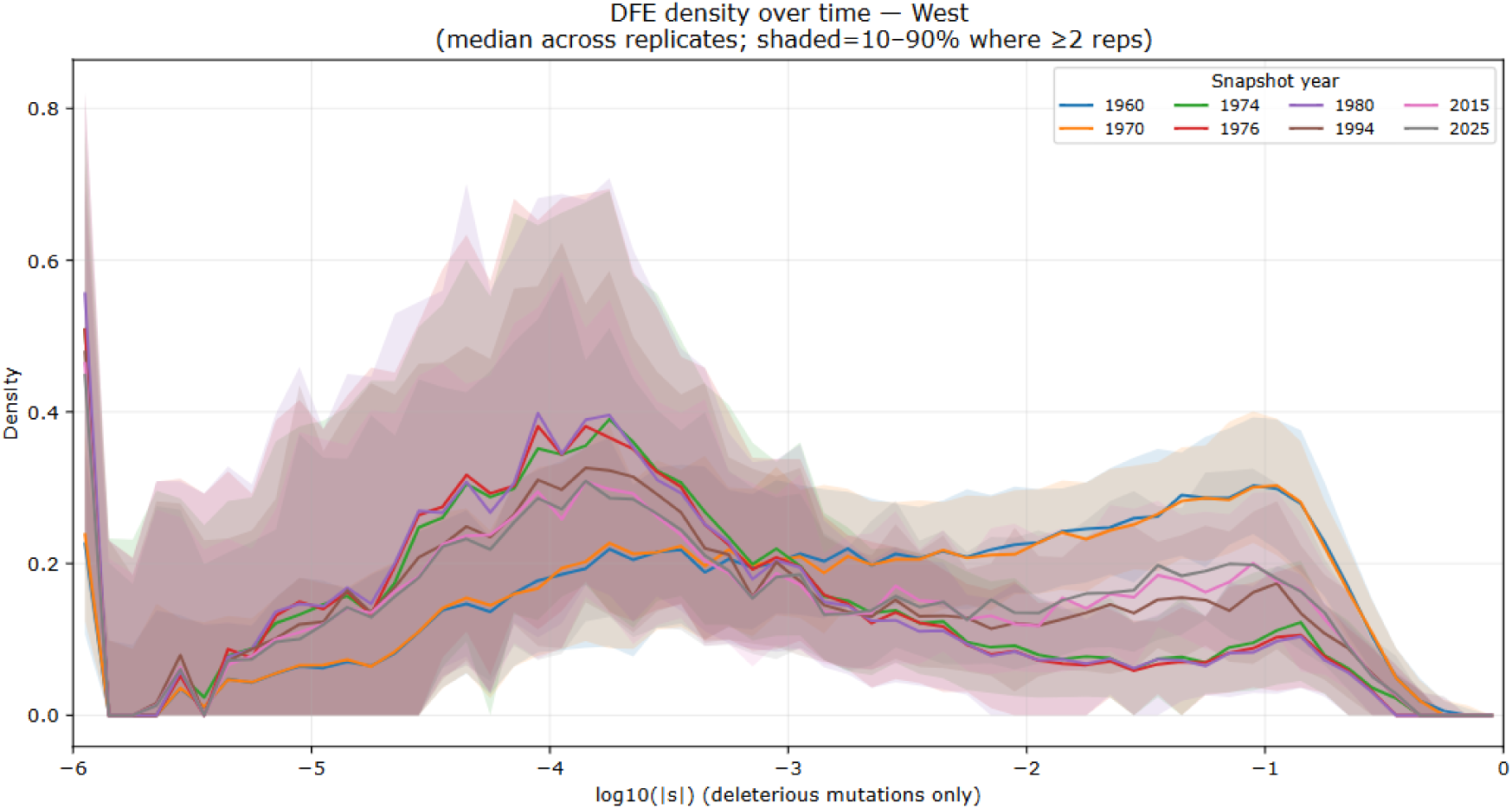
Changes of standing DFE through time in simulations. The shift of DFE of standing deleterious mutations towards a lower selection coefficient in simulations from 1960 to 1980, and an increase of density of highly deleterious mutations after 1994.

