## Supplementary Information for "Conservation rescued the Mauritius kestrel from extinction but not from genomic erosion"

**The PDF file includes:**

Supplementary Methods

Supplementary Figs. 1-7

Supplementary Tables 1-5

References

**Supplementary Methods**

**Simulated populations with SLiM**

Individuals were diploid, autosomal and sex was assigned. The model included two focal subpopulations corresponding to the western remnant population in Black River Gorges and the eastern reintroduced population in the Bambou Mountains. The west population was present throughout the simulation. The east population was founded from west individuals in the first year for which the empirical east population trajectory was non-zero. After founding, the west and east populations followed separate annual target trajectories. No routine natural migration was included after the founding of the east population^1^.

**Simulated genome architecture**

Rather than simulating the full kestrel genome which requires intensive computing resources, we used a reduced genomic representation. The simulated genome comprised two components. The first was a chromosome-scale region based on the collared flycatcher chromosome 12 feature map and recombination map^2–4^. The second was an appended exome-like component consisting of 18,000 genes of 1,500 bp each, distributed across 20 chromosomes. Together, the simulated genomic representation was approximately 48 Mb and accurately represented linkage dynamics. Recombination in the chromosome-scale region followed the input recombination map. In the appended exome-like component, recombination was set to zero within genes and to 1×10^-4^ between adjacent genes with chromosome boundaries assigned a recombination probability of 0.5 to represent independent segregation. This avoided treating the compacted exome-like component as a single fully linked region.

Deleterious mutations were assigned according to the genomic region type. Relative to neutral mutations, deleterious mutations were enriched in exonic regions, occurred at lower relative probability in intronic regions and were absent from intergenic regions. Specifically, deleterious-to-neutral mutation weights were 2.31 in exons, 0.8 in introns, and 0 in intergenic regions. The appended exome-like component used exon weighting^5^.

The per-site mutation rate was set to 5.0 x 10^-9^ per site per generation. Because mutation rate and ancestral effective size jointly influence equilibrium diversity, we treated the mutation rate as uncertain and evaluated appropriate values via sensitivity analyses. Alternative mutation rates shifted the absolute level of simulated heterozygosity, as expected, but did not alter the qualitative post-bottleneck genomic trajectory: heterozygosity declined, F_ROH_ increased, masked load declined and realised load increased or stabilised depending on the time period (Supplementary Fig. 4).

**Distribution of deleterious fitness effects**

Deleterious selection coefficients were drawn from a gamma distribution with a tail of lethal mutations, following conservation-genomic simulation approaches that model both weakly deleterious and strongly deleterious mutations^6^. The main distribution consisted of gamma-distributed negative selection coefficients and a recessive lethal class following Kardos et al. 2021 ^7^. Because simulation ticks represented years, selection coefficients were applied as annual viability effects in the non-Wright-Fisher life cycle.

**Burn-in and ancestral census-size calibration**

The ancestral census prior was calibrated against the empirical historical genomic baseline rather than treated as an independently estimated demographic parameter. We compared alternative ancestral census-size ranges and retained the range that reproduced the low but non-zero heterozygosity observed in the historical samples before the recent bottleneck. Smaller ancestral census priors produced too little standing heterozygosity and therefore failed to recover the empirical heterozygosity trajectory before and after the bottleneck (Supplementary Fig. 5). We used an ancestral census size of 6,000-12,000 because it provided a reasonable match to the empirical heterozygosity. This parameter was used to set the starting genomic condition for the forward simulations, not as an estimate of long-term effective population size or historical abundance.

The final simulations used a 40,000-year burn-in before imposing the historical decline. We tested burn-in lengths of 20,000, 40,000, 80,000 and 100,000 years (Supplementary Fig. 5). Post-bottleneck trajectories of heterozygosity, F_ROH_, deleterious allele burden and load indices were similar across burn-in lengths (Supplementary Fig. 6), indicating that the main conclusions were not sensitive to burn-in duration. The 40,000 year burn-in was therefore used for the main analyses as a compromise between mutation-selection-drift equilibrium and computational tractability.

**Historical demography and the bottleneck in 1974**

After the burn-in stage, the simulated west population declined from the ancestral census size toward the historical bottleneck using an exponential decline trajectory beginning in 1500. This decline represented long-term habitat degradation before the documented 20th century crash^1,8^. From the recovery period onward, empirical demographic trajectories were used as annual target census sizes for west and east populations. Annual target sizes were used to guide density-dependent regulation rather than to set population size directly. In each year, the target size is used to scale the fitness measure related to survival, but the realised number of individuals remained free to vary because reproduction, mortality, genetic fitness, demographic stochasticity and scenario-specific management effects were all simulated explicitly. A four-individual bottleneck was imposed in 1974, reflecting the documented crash to four known wild birds ^9^. We tested for alternative bottleneck sizes of 6, 8, and 10, and lifted bottleneck sizes yielded lower F_ROH_ than empirical results (Supplementary Fig. 7). The bottleneck was implemented by retaining four individuals and setting survival of all other west individuals to zero in that year. Retained individuals were preferentially sampled from breeding-age birds and where possible sex-balanced. This preserved the genomic severity of a four-individual bottleneck while avoiding artefactual extinction caused by retaining only senescent individuals or only one sex.

**Life history, mating system and reproduction**

Age-specific mortality was read from a Mauritius kestrel life table. Individuals entered the breeding pool at age 1 and could breed until age 12. The oldest age class was assigned a mortality probability of 1.0 so that lifespan remained bounded by the age range represented in the life table.

Reproduction was sex-specific and monogamous. Eligible breeders were males and females within the breeding-age range. Existing male-female pairs remained paired across years unless one member died. Unpaired eligible males and females were randomly paired within each subpopulation. Each pair had a stage-specific probability of breeding. Conditional on breeding, annual reproductive output was drawn from a gamma-Poisson distribution with mean parameter 3 and shape parameter 40 and was capped at five offspring per breeding event. Offspring entered the population as age-0 individuals and were unpaired until reaching breeding age.

**Density regulation and carrying capacity**

Density regulation acted through survival rather than by directly setting population size. For each population and year, the empirical demographic trajectory provided a target abundance used to guide density-dependent survival scaling. Survival was reduced when the expected number of survivors exceeded this target but was not increased above 1.0 when the population was below target. Realised abundance could therefore deviate from the empirical trajectory because reproduction, mortality, demographic stochasticity, releases and scenario-specific management effects were simulated explicitly^10–12^.

**Historical management stages**

Historical changes in environmental conditions and conservation management were represented by modifying age-specific mortality and pair breeding probability across broad demographic stages (Supplementary Table 4). These values were used as process-model parameters not as independent field estimates of every ecological mechanism. They allowed the model to represent the timing of long-term habitat degradation, the crash period, intensive management and later supportive management while remaining anchored to the empirical demographic trajectories^10–12^.

**Captive releases and reintroduction**

Historical releases were included in the baseline and conservation-stops scenarios. We modelled year-specific releases into the western and eastern populations between 1984 and 1993, matching the reported totals^10^ for the two focal populations. We did not model a separate captive pedigree due to the lack of records. Instead, released juveniles were generated by crossing randomly sampled opposite-sex individuals from the western source population and adding the resulting age-0 offspring to the target population. This approximated the genomic contribution of the captive-breeding and release programme while retaining the western remnant population as the source of released genetic variation. Released individuals subsequently experienced the same survival, genetic fitness, density regulation and breeding processes as wild-born individuals.

**Management scenarios**

We simulated three management scenarios. Firstly, the ‘baseline scenario’ followed the historical decline, the 1974 bottleneck, documented captive releases, eastern founding, empirical recovery trajectories and continued supportive management into the future (Supplementary Table 4). Secondly, the ‘no-conservation scenario’ represented a counterfactual in which conservation management was not implemented after the crash in 1974. In this scenario, post-crash survival and breeding remained poor and conservation actions such as captive releases and eastern reintroduction were disabled. This scenario tested whether the species could recover demographically without management after the bottleneck. Thirdly, the ‘conservation-stops scenario’ followed the baseline historical trajectory until 2025. After 2025, supportive management was removed by reducing breeding probability and increasing mortality. For each scenario, we ran 100 replicate simulations and recorded the number of simulations that resulted in extinction. Extinction was defined as the loss of all individuals across the simulated focal metapopulation.

**Simulation outputs**

For all simulation-derived genomic summaries, metrics were calculated at the individual level where applicable and then averaged across sampled individuals within each output year, population and replicate. When west and east populations were combined, summaries were calculated across all sampled individuals from the focal metapopulation for that year and replicated. Plotted trajectories were then generated by calculating the median across simulation replicates at each time point; uncertainty around trajectories is shown as the 10-90% range unless otherwise stated.

**Model calibration and empirical checking**

We compared simulated trajectories with the empirical temporal genomic data before interpreting counterfactual or future scenarios. The empirical dataset contained discrete sampling points spanning historical specimens, early post-bottleneck individuals, mid-recovery individuals and recent individuals. Simulations were therefore used to infer continuous annual trajectories between empirical time points. We therefore used the empirical genomic data as calibration and model-checking targets, not merely as post hoc comparisons.

**Simulation-anchored weighted load partitioning**

SnpEff impact classes and CADD scores provide useful annotations on the function or deleteriousness of the variants, but raw class counts cannot be used directly to estimate realised load, masked load or purging because the classes are not additive on a fitness scale. For example, one LOF variant is not necessarily equivalent to two missense variants. Furthermore, the annotations have no direct lethal-equivalent interpretation or estimated fitness impact. Therefore, simple counts of variants can show whether putatively deleterious variants change through time, but they cannot by themselves quantify how much masked load has been converted into realised load, or whether total fitness-weighted load has declined through purging.

To address this, we developed an exploratory weighted load-partitioning analysis that converts various annotation classes into approximate lethal-equivalent units using simulation-anchored class-specific coefficients. This analysis should be interpreted as a calibrated load-partitioning approach rather than as a definitive estimate of the true selection coefficient of every variant in each annotation class. The objective was to use variant-impact classes as effect-weighted proxies for genetic load and to test whether the temporal genomic data show the expected signatures of masked-to-realised load conversion and purging after demographic recovery. We assumed recessive effects for this exploratory analysis, so heterozygous variants contribute to masked load but not realised load. For individual *i*, realised load (RL), masked load (ML) and total genetic load (GL) were calculated as:

*GL_i = RL_i + ML_i*

SnpEff:

*RL_i = s_HIGH x HIGH_hom,i + s_MODERATE x MODERATE_hom,i*

*ML_i = 0.5 x (s_HIGH x HIGH_het,i + s_MODERATE x MODERATE_het,i)*

CADD:

*RL_i = s_CADD_01 x CADD_01_hom,i + s_CADD_05 x CADD_05_hom,i*

*ML_i = 0.5 x (s_CADD_01 x CADD_01_het,i + s_CADD_05 x CADD_05_het,i)*

Here, HIGH_hom and MODERATE_hom are the corrected counts of homozygous HIGH and MODERATE SnpEff variants; CADD_01_hom and CADD_05_hom are the corrected counts of homozygous variants with top 1% and top 5% CADD score; HIGH_het and MODERATE_het are the corresponding corrected heterozygous counts; and CADD_01_het and CADD_05_het are the corrected counts of heterozygous variants with top 1% and top 5% CADD score. To correct for variant-calling quality, the corrected counts were calculated by dividing the raw counts of each individual by the number of derived neutral alleles, and then multiplied by the average number of derived neutral alleles. LOW-impact variants and variants with CADD below top 5% were excluded from the main analysis because previous exploratory fitting indicated that these classes were not separately identifiable from higher classes in these data. We calibrated the coefficients using the post-bottleneck temporal genomic sample (n = 59: early post-bottleneck n = 23, middle n = 9, recent n = 27) and the simulation-derived mean genetic-load trajectory. We fitted linear model with unconstrained least-squares optimization in R:

SnpEff:

*mean_GL_group = s_HIGH x (HIGH_hom + 0.5 x HIGH_het) + s_MODERATE x (MODERATE_hom + 0.5 x MODERATE_het)*

CADD:

*mean_GL_group = s_CADD_01 x (CADD_01_hom + 0.5 x CADD_01_het) + s_CADD_05 x (CADD_05_hom + 0.5 x CADD_05_het)*

We used group-level simulation targets: early post-bottleneck samples were matched to the mean simulated GL across 1987-1995, middle samples to the 2011 simulated GL value, and recent samples to the mean simulated GL across 2019-2021. We assessed sensitivity by repeating the simulation-anchored calibration after omitting each post-bottleneck individual in turn (Supplementary Table 5a). We further tested with raw counts of variants and had similar estimations (Supplementary Table 5b). Historical samples were excluded from the coefficient calibration to avoid issues with lower-quality data and potential shift of standing DFE (Fig. 2G).

**Supplementary Figures**

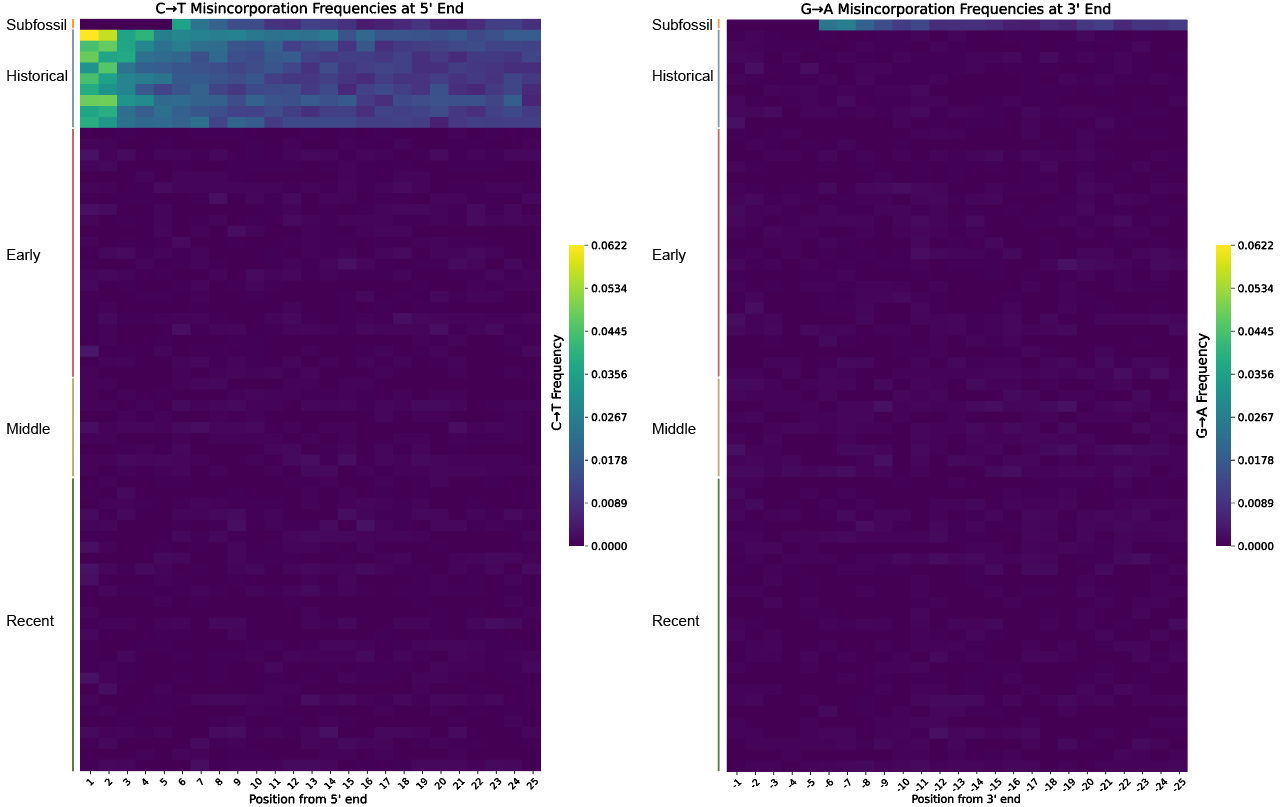

**Supplementary Fig. 1** **Misincorporation frequencies on both ends of the reads for each Mauritius kestrel sample inferred with MapDamage.**

Each row represents one sample and each column represents one bp position on the read. The color of the cell represents the C to T or G to A transition rate. The first and last five base pairs of the reads from the subfossil sample were masked and thus had no misincorporation frequencies for those positions.

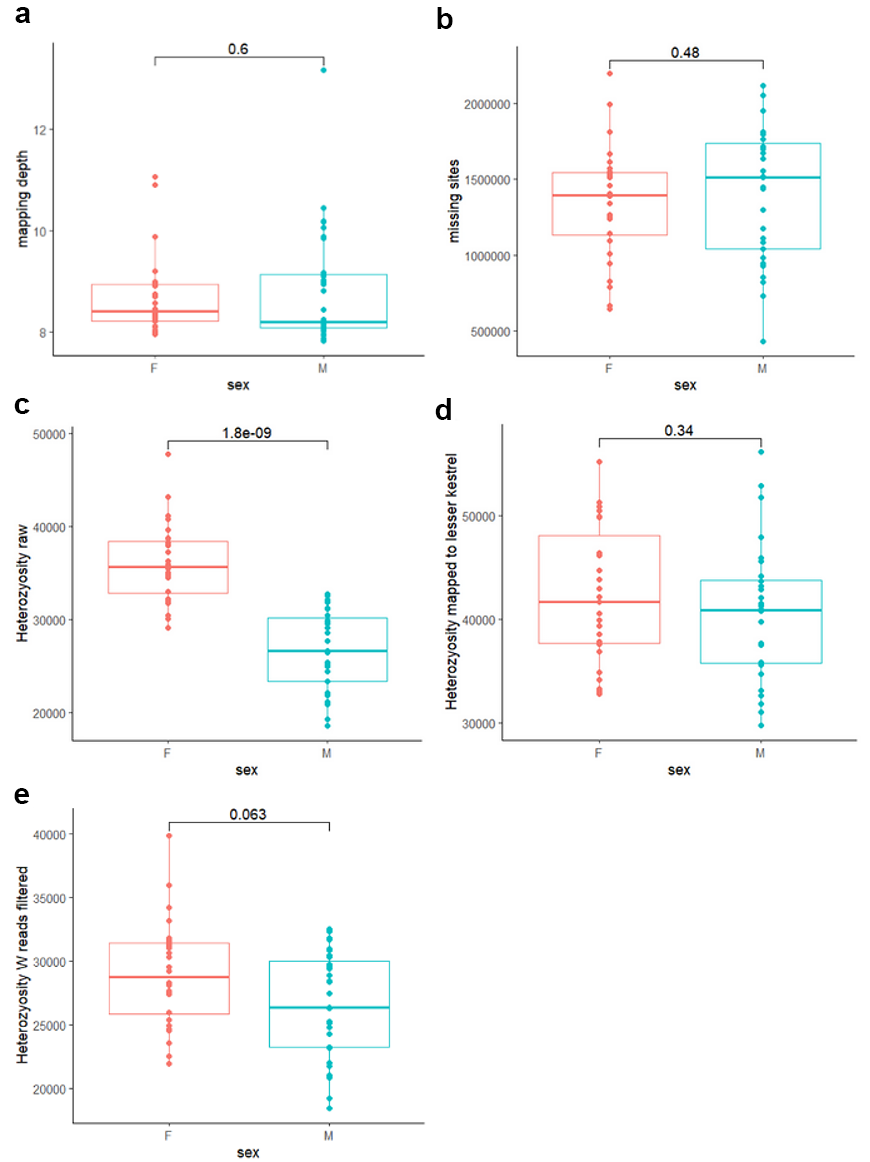

**Supplementary Fig. 2** **Mapping bias due to the lack of W chromosome in the reference genome of Mauritius kestrel.**

The mapping depth (**a**) and number of missing sites (**b**) had no difference between the sexes, and the number of heterozygous sites was not significantly higher in females (F) than in males (M) when mapped to the reference genome of the lesser kestrel (**c**) which has the W chromosome. Thus, the higher heterozygosity shown in females when mapped to the reference genome of the Mauritius kestrel (**d**) was likely an artifact due to the lack of W chromosome. **e**, When excluding the reads mapped to the W chromosome of the lesser kestrel, such bias was corrected.

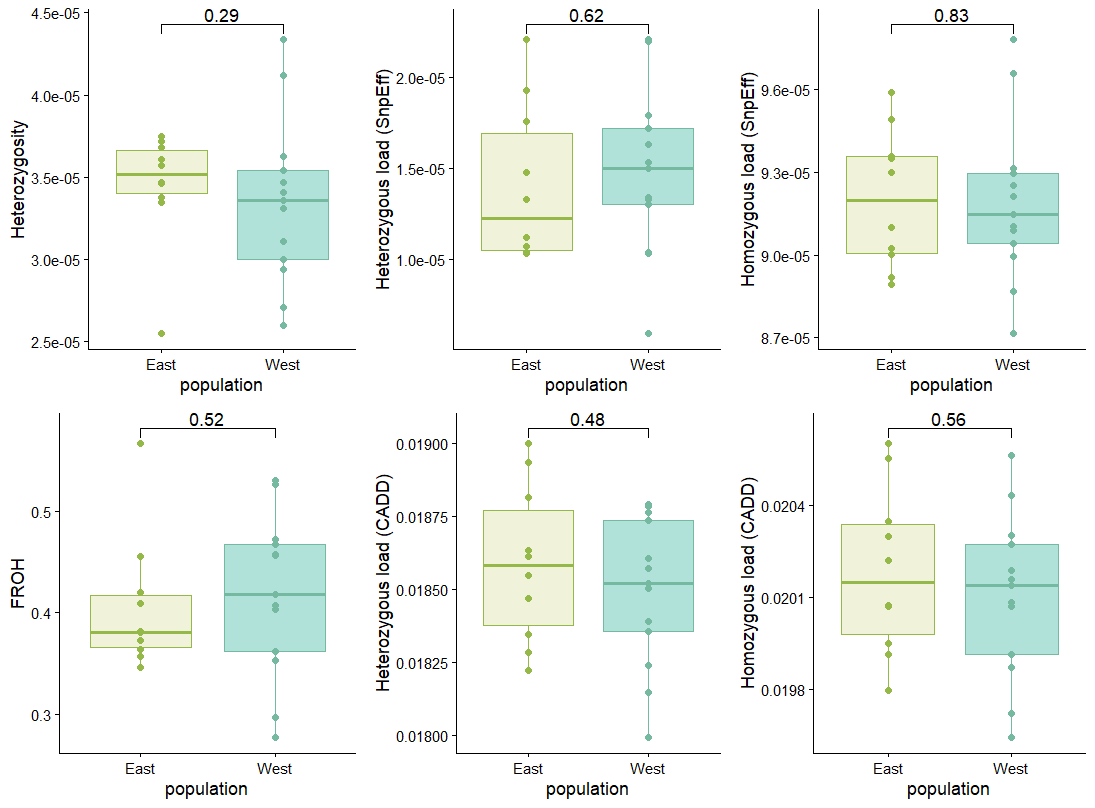

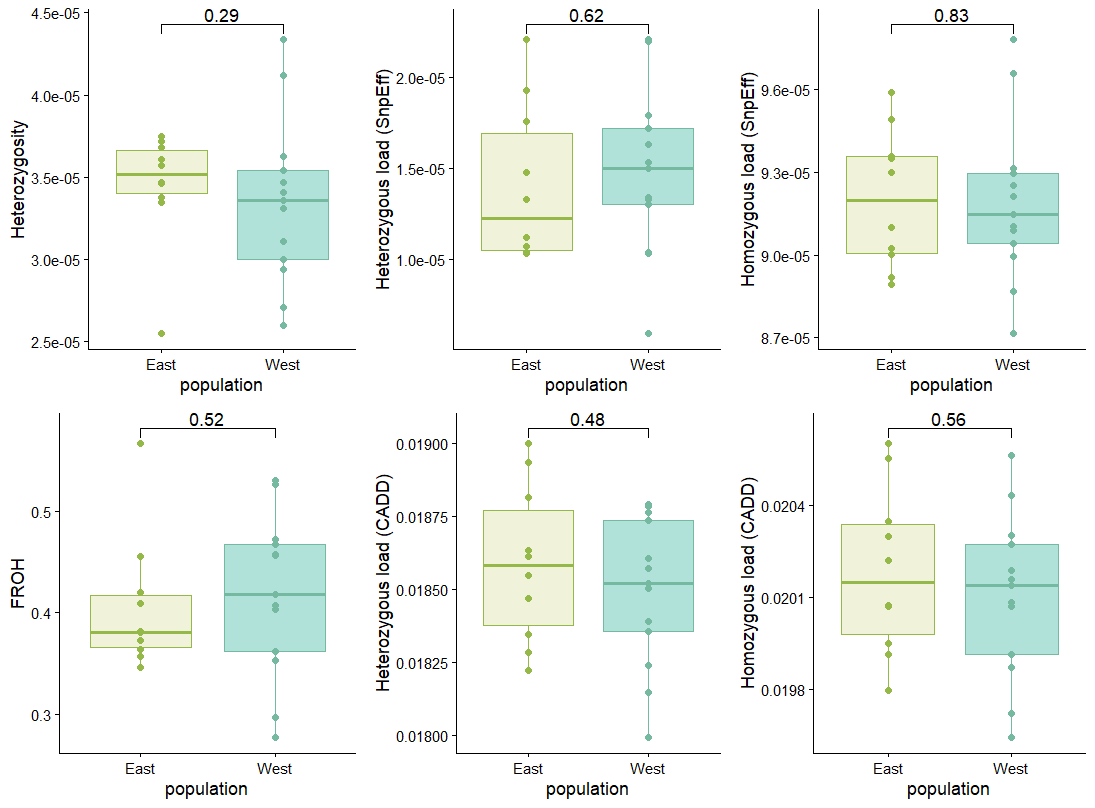

**Supplementary Fig. 3 Comparison of genetic diversity and inbreeding of two populations in the 1990s.**

Both heterozygosity and F_ROH_ showed no significant difference between the two populations.

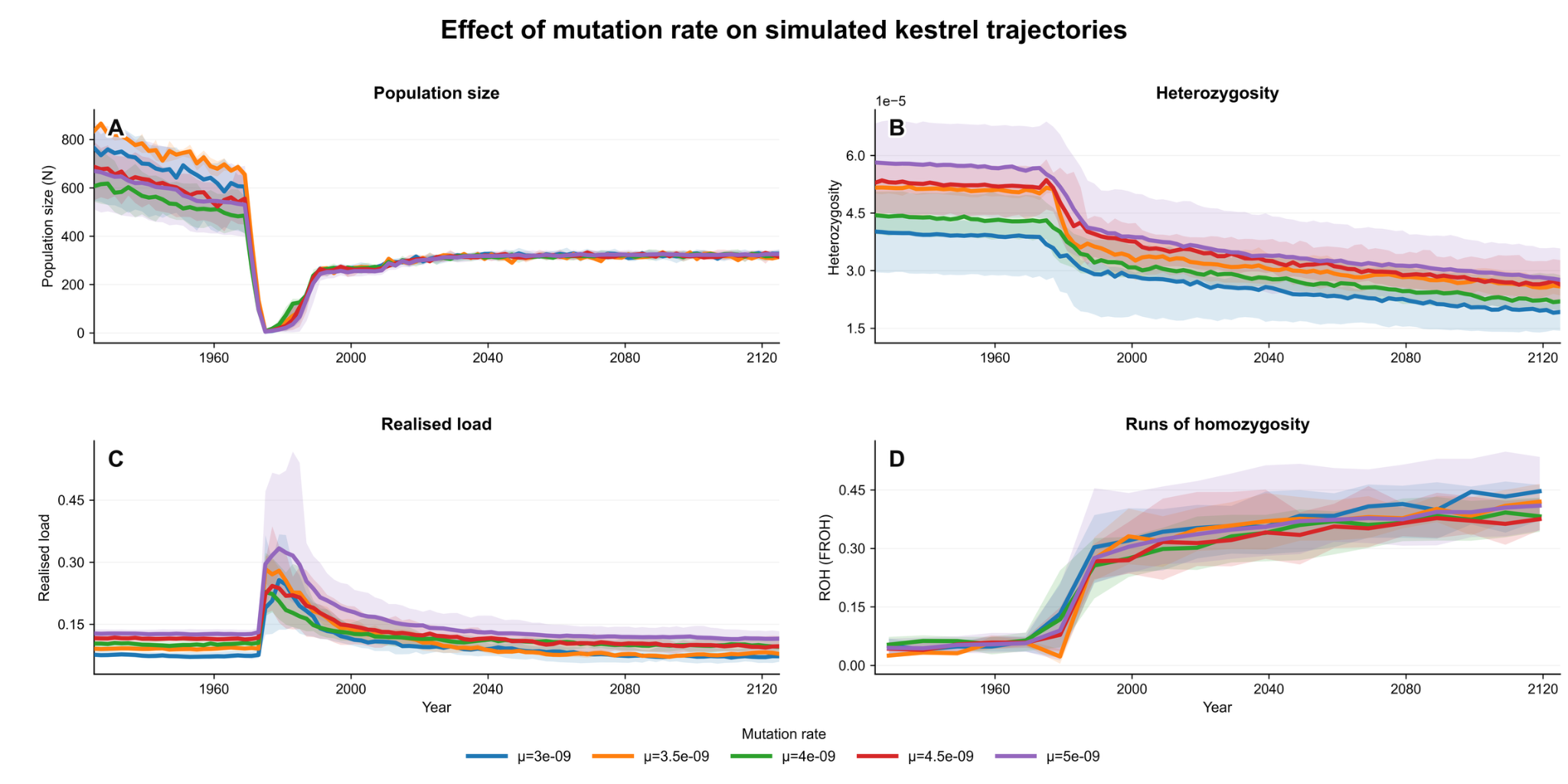

**Supplementary Fig. 4 Various mutation rates tested in SLiM simulations showing the population and genetic trends are consistent.**

For each mutation rate, 25 replicates were performed. Solid lines represent the mean value and the ribbons represent the 10-90% range.

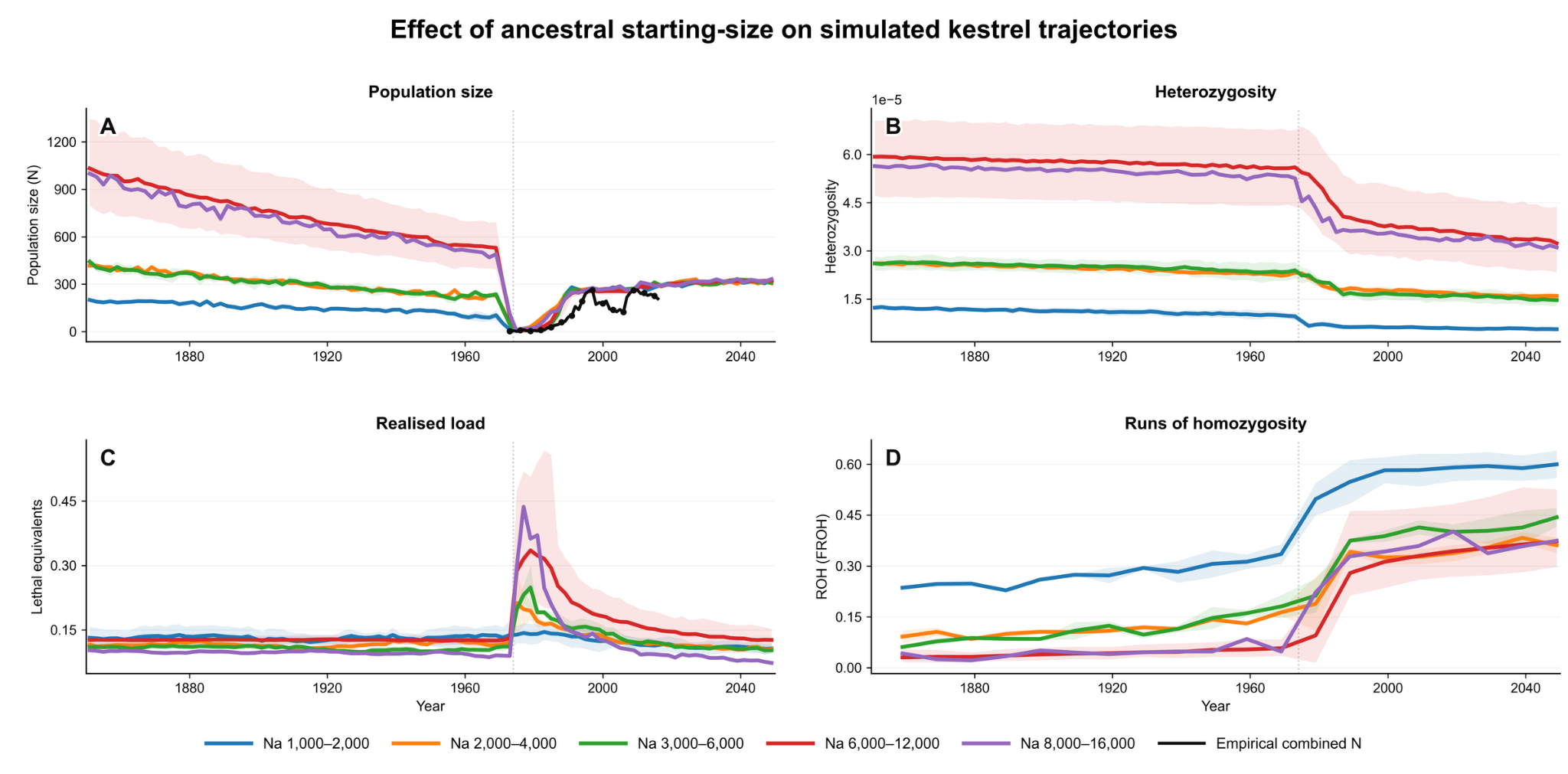
**Supplementary Fig. 5 Testing of effects of ancestral population size on demographic changes, heterozygosity, genetic load and F_ROH_.**
For each ancestral population size, 25 replicates were performed. Solid lines represent the mean value and the ribbons represent the 10-90% range.

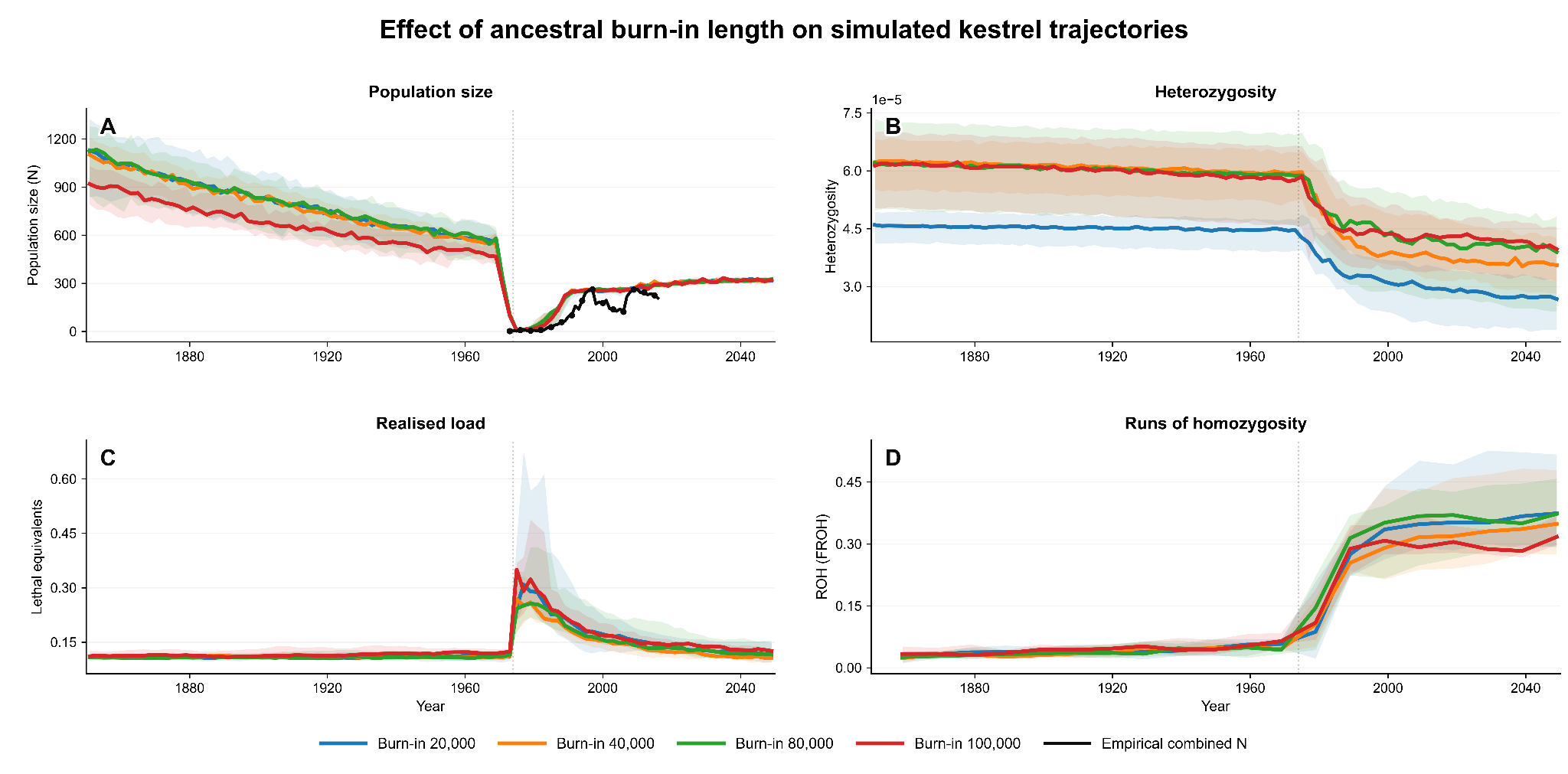

**Supplementary Fig. 6 Testing of effects of changing the length of burn-in period on demographic changes, heterozygosity, genetic load and F_ROH_.**

For each burn-in period value, 25 replicates were performed. Solid lines represent the mean value and the ribbons represent the 10-90% range.

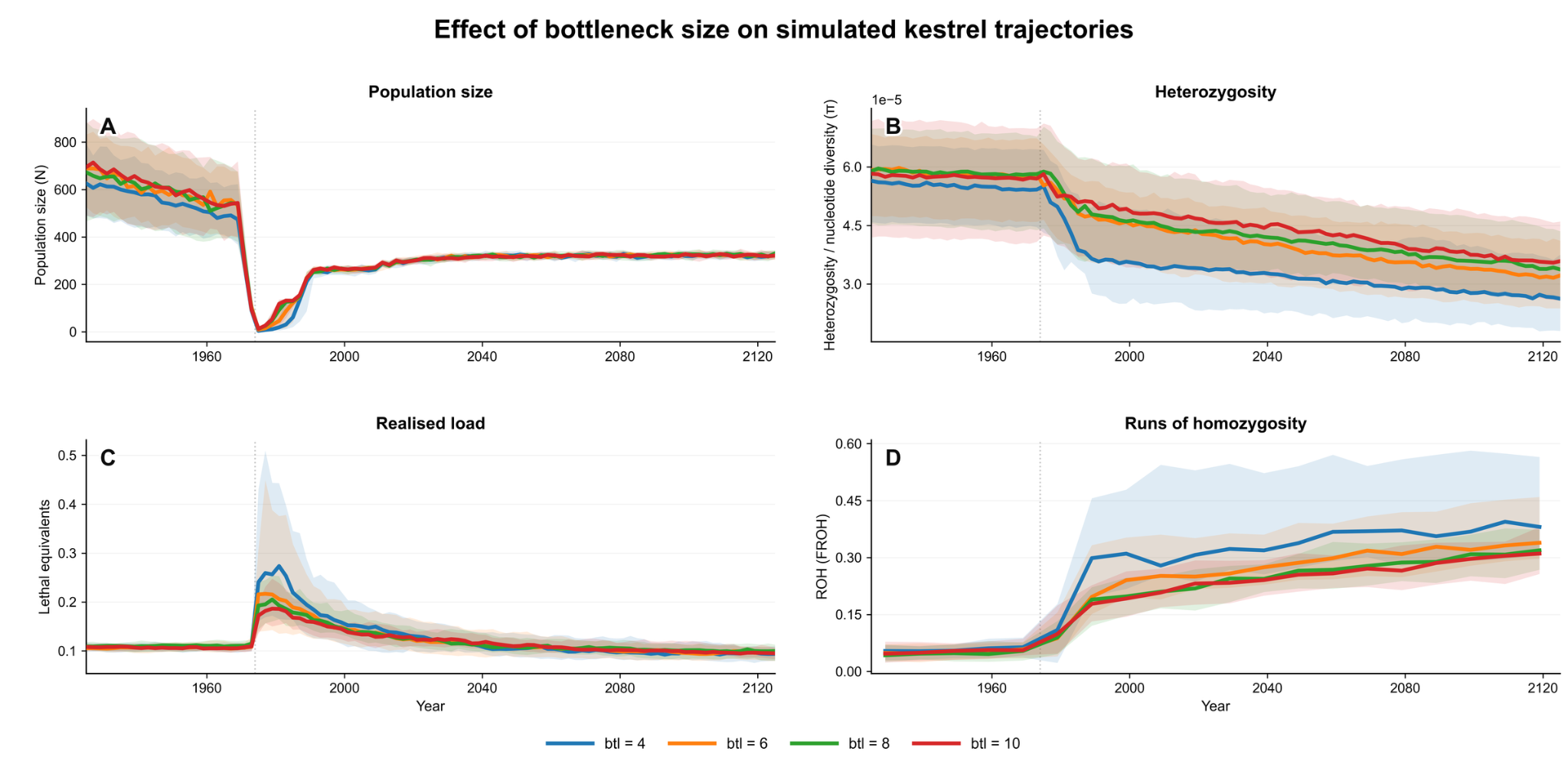

**Supplementary Fig. 7 Changing the population size at the bottleneck from 4 to 6, 8 or 10 in SLiM simulations did not change population and genetic trends.**

For each bottleneck size, 25 replicates were performed. Solid lines represent the mean value and the ribbons represent the 10-90% range.

**Supplementary Tables**

**Supplementary Table 1**. Samples of Mauritius kestrels for temporal comparisons.

| **Sample** | **Time period** | **Sex** | **Population** | **Location** | **Year** | **Depth** | **Note** | **Accession** |
| --- | --- | --- | --- | --- | --- | --- | --- | --- |
| aMK_SB787 |  | M |  | NHMUK | 1769±17 | 8.7 | subfossil bone | SRRxxxxxxxx |
| hMK_1 | Historical | F |  | NHMUK | <1843 | 8.7 |  | SRRxxxxxxxx |
| hMK_10 | Historical | F |  | NHMUK | 1859-1867 | 6.1 |  | SRRxxxxxxxx |
| hMK_11 | Historical | F |  | MNHN | 1834 | 8.0 |  | SRRxxxxxxxx |
| hMK_15 | Historical | M |  | UMZC | 1864 | 2.1 | only mtDNA | SRRxxxxxxxx |
| hMK_16 | Historical | M |  | UMZC | 1864 | 8.6 |  | SRRxxxxxxxx |
| hMK_2 | Historical | M |  | NHMUK | 1866 | 6.1 |  | SRRxxxxxxxx |
| hMK_20 | Historical | M |  | UMZC | 1860 | 11.6 |  | SRRxxxxxxxx |
| hMK_22 | Historical | M |  | UMZC | 1866 | 12.1 |  | SRRxxxxxxxx |
| hMK_23 | Historical | F |  | UMZC | 1832 | 16.6 |  | SRRxxxxxxxx |
| hMK_25 | Historical | M |  | MCZ | N/A | 8.4 |  | SRRxxxxxxxx |
| hMK_26 | Historical | M |  | MCZ | N/A | 4.7 | only mtDNA | SRRxxxxxxxx |
| hMK_4 | Historical | M |  | Leiden | N/A | 4.4 | only mtDNA | SRRxxxxxxxx |
| hMK_5 | Historical | F |  | Leiden | 1875 | 4.5 | only mtDNA | SRRxxxxxxxx |
| hMK_7 | Historical | F |  | NHMUK | 1858 | 3.9 | only mtDNA | SRRxxxxxxxx |
| hMK_8 | Historical | M |  | NHMUK | N/A | 3.2 | only mtDNA | SRRxxxxxxxx |
| MK_413 | Early | M | West | West Cliffs | 1995 | 13.2 | downsampled | SRRxxxxxxxx |
| MK_421 | Early | M | West | Acacia Rock | 1993 | 8.0 | downsampled | SRRxxxxxxxx |
| MK_43 | Early | F | West | Brise Fer Bluff | 1996 | 11.1 | downsampled | SRRxxxxxxxx |
| MK_446 | Early | M | West | Caves | 1993 | 8.9 | downsampled | SRRxxxxxxxx |
| MK_591782 | Early | M | West | Zaco | 1993 | 10.1 | downsampled | SRRxxxxxxxx |
| MK_591798 | Early | F | West | Caves | 1994 | 9.2 | downsampled | SRRxxxxxxxx |
| MK_594332 | Early | F | East | Ferney 5 | 1994 | 8.0 | downsampled | SRRxxxxxxxx |
| MK_594337 | Early | F | East | Ylang Ylang | 1994 | 9.0 | downsampled | SRRxxxxxxxx |
| MK_594338 | Early | F | East | Ylang Ylang | 1994 | 8.3 | downsampled | SRRxxxxxxxx |
| MK_594341 | Early | M | East | Mt Camizard 1 | 1994 | 9.9 | downsampled | SRRxxxxxxxx |
| MK_594342 | Early | M | East | Mt Camizard 1 | 1995 | 8.4 | downsampled | SRRxxxxxxxx |
| MK_594345 | Early | F | East | Ferney 7 | 1994 | 8.6 | downsampled | SRRxxxxxxxx |
| MK_594350 | Early | F | East | Mt Camizard 2 | 1994 | 8.3 | downsampled | SRRxxxxxxxx |
| MK_594356 | Early | F | West | Eden Flycatcher | 1994 | 8.9 | downsampled | SRRxxxxxxxx |
| MK_594357 | Early | M | West | Morne Seche Caves | 1994 | 7.8 | downsampled | SRRxxxxxxxx |
| MK_594358 | Early | M | West | Eden Flycatcher | 1994 | 8.0 | downsampled | SRRxxxxxxxx |
| MK_594369 | Early | M | West | Stoneman | 1994 | 9.2 | downsampled | SRRxxxxxxxx |
| MK_594370 | Early | F | West | Rempart Ridge | 1994 | 9.9 | downsampled | SRRxxxxxxxx |
| MK_594371 | Early | F | West | Tamarin Tree | 1994 | 8.5 | downsampled | SRRxxxxxxxx |
| MK_594373 | Early | M | West | Le Morne | 1994 | 10.4 | downsampled | SRRxxxxxxxx |
| MK_594387 | Early | M | East | Etoile 9 | 1995 | 8.1 | downsampled | SRRxxxxxxxx |
| MK_596254 | Early | F | East | Le Vallon Power Station Pair | 1996 | 8.9 |  | SRRxxxxxxxx |
| MK_596306 | Middle | M | East | Mt Camizard 1 | 1995 | 8.1 |  | SRRxxxxxxxx |
| MK_5A12345 | Middle | M | East | Re11 | 2009 | 8.8 |  | SRRxxxxxxxx |
| MK_5A12347 | Middle | M | East | Re11 | 2009 | 7.9 |  | SRRxxxxxxxx |
| MK_5A12512 | Middle | F | East | Fe4 | 2009 | 8.3 |  | SRRxxxxxxxx |
| MK_5A12514 | Middle | M | East | Fe8 | 2009 | 8.2 |  | SRRxxxxxxxx |
| MK_5A12515 | Middle | F | East | Fe8 | 2009 | 9.0 |  | SRRxxxxxxxx |
| MK_5A12527 | Middle | M | East | YY | 2009 | 8.2 |  | SRRxxxxxxxx |
| MK_5A12528 | Middle | M | East | YY | 2009 | 9.0 |  | SRRxxxxxxxx |
| MK_5A12529 | Middle | F | East | MC5 | 2009 | 8.0 |  | SRRxxxxxxxx |
| MK_5A12533 | Middle | F | East | ET5 | 2009 | 8.8 |  | SRRxxxxxxxx |
| MK_5A19552 | Recent | M | East | Ferney Enclosure | 2019 | 8.2 |  | SRRxxxxxxxx |
| MK_5A19559 | Recent | M | East | Ferney Enclosure | 2019 | 10.2 |  | SRRxxxxxxxx |
| MK_5A19569 | Recent | F | East | Le Vallon 6 | 2019 | 8.7 |  | SRRxxxxxxxx |
| MK_5A19603 | Recent | F | East | Ferney 3 | 2020 | 8.9 |  | SRRxxxxxxxx |
| MK_5A19607 | Recent | F | East | Domaine du Chasseur Cabin | 2020 | 8.0 |  | SRRxxxxxxxx |
| MK_5A19608 | Recent | F | East | Ferney Ingam | 2020 | 8.0 |  | SRRxxxxxxxx |
| MK_5A19609 | Recent | M | East | Domaine de l'Etoile 5 | 2020 | 8.2 |  | SRRxxxxxxxx |
| MK_5A19613 | Recent | F | East | Ferney 10 | 2020 | 8.2 |  | SRRxxxxxxxx |
| MK_5A19614 | Recent | F | East | Ferney Ingam | 2020 | 8.9 |  | SRRxxxxxxxx |
| MK_5A19615 | Recent | M | East | Mt Camizard 5 | 2021 | 10.2 |  | SRRxxxxxxxx |
| MK_5A19617 | Recent | M | East | Ferney Middle | 2020 | 8.0 |  | SRRxxxxxxxx |
| MK_5A19624 | Recent | M | East | Ferney Middle | 2021 | 8.2 |  | SRRxxxxxxxx |
| MK_5A19933 | Recent | M | East | Ferney 11 | 2019 | 9.1 |  | SRRxxxxxxxx |
| MK_5A19951 | Recent | F | East | Petit Parc | 2019 | 8.4 |  | SRRxxxxxxxx |
| MK_5A19960 | Recent | M | East | Ferney 3 | 2020 | 8.2 |  | SRRxxxxxxxx |
| MK_5A19963 | Recent | F | East | Ferney 11 | 2019 | 8.3 |  | SRRxxxxxxxx |
| MK_5A19966 | Recent | F | East | Riche en Eau 3 | 2019 | 8.1 |  | SRRxxxxxxxx |
| MK_5A19969 | Recent | M | East | Ferney Cliff | 2019 | 8.0 |  | SRRxxxxxxxx |
| MK_5A19992 | Recent | M | East | Riche en Eau 3 | 2020 | 8.1 |  | SRRxxxxxxxx |
| MK_5A19996 | Recent | M | East | Riche en Eau 3 | 2020 | 9.8 |  | SRRxxxxxxxx |
| MK_5A19997 | Recent | F | East | Power Station Cliff | 2020 | 8.1 |  | SRRxxxxxxxx |
| MK_5A26603 | Recent | M | East | Petit Parc | 2021 | 9.0 |  | SRRxxxxxxxx |
| MK_5A26607 | Recent | M | East | Ferney 3 | 2021 | 8.2 |  | SRRxxxxxxxx |
| MK_5A26610 | Recent | F | East | Ferney Dupont | 2021 | 8.2 |  | SRRxxxxxxxx |
| MK_5A26612 | Recent | F | East | Domaine du Chasseur 8 | 2021 | 8.3 |  | SRRxxxxxxxx |
| MK_5A26613 | Recent | M | East | Domaine du Chasseur 8 | 2021 | 7.9 |  | SRRxxxxxxxx |
| MK_5A26614 | Recent | F | East | Domaine du Chasseur 8 | 2021 | 10.9 |  | SRRxxxxxxxx |

**Supplementary Table 2**. *Falco* kestrel samples for interspecific comparisons.

| Sample | Common name | Species | Sex | Depth | Accession |
| --- | --- | --- | --- | --- | --- |
| CanK_C5971 | Canary Islands kestrel | Falco tinnunculus canariensis | M | 19.9 | SRRxxxxxxxx |
| CanK_C5973 | Canary Islands kestrel | Falco tinnunculus canariensis | F | 24.0 | SRRxxxxxxxx |
| CanK_C5975 | Canary Islands kestrel | Falco tinnunculus canariensis | F | 19.4 | SRRxxxxxxxx |
| CanK_C5976 | Canary Islands kestrel | Falco tinnunculus canariensis | F | 17.3 | SRRxxxxxxxx |
| CanK_C5977 | Canary Islands kestrel | Falco tinnunculus canariensis | F | 29.1 | SRRxxxxxxxx |
| CanK_C5978 | Canary Islands kestrel | Falco tinnunculus canariensis | F | 31.0 | SRRxxxxxxxx |
| GK1 | Greater kestrel | Falco rupicoloides | M | 19.4 | SRRxxxxxxxx |
| GK10 | Greater kestrel | Falco rupicoloides | M | 22.8 | SRRxxxxxxxx |
| GK2 | Greater kestrel | Falco rupicoloides | M | 20.3 | SRRxxxxxxxx |
| GK3 | Greater kestrel | Falco rupicoloides | M | 16.3 | SRRxxxxxxxx |
| GK4 | Greater kestrel | Falco rupicoloides | F | 16.4 | SRRxxxxxxxx |
| GK5 | Greater kestrel | Falco rupicoloides | F | 17.7 | SRRxxxxxxxx |
| GK6 | Greater kestrel | Falco rupicoloides | F | 15.7 | SRRxxxxxxxx |
| GK7 | Greater kestrel | Falco rupicoloides | M | 18.6 | SRRxxxxxxxx |
| GK8 | Greater kestrel | Falco rupicoloides | F | 17.7 | SRRxxxxxxxx |
| GK9 | Greater kestrel | Falco rupicoloides | F | 17.2 | SRRxxxxxxxx |
| LK1 | Lesser kestrel | Falco naumanni | M | 19.7 | SRRxxxxxxxx |
| LK2 | Lesser kestrel | Falco naumanni | M | 16.5 | SRRxxxxxxxx |
| LK3 | Lesser kestrel | Falco naumanni | F | 19.3 | SRRxxxxxxxx |
| LK4 | Lesser kestrel | Falco naumanni | F | 20.7 | SRRxxxxxxxx |
| LK5 | Lesser kestrel | Falco naumanni | F | 19.5 | SRRxxxxxxxx |
| LK6 | Lesser kestrel | Falco naumanni | M | 16.6 | SRRxxxxxxxx |
| LK7 | Lesser kestrel | Falco naumanni | M | 17.5 | SRRxxxxxxxx |
| LK8 | Lesser kestrel | Falco naumanni | M | 16.0 | SRRxxxxxxxx |
| MadKes1 | Madagascar kestrel | Falco newtoni | M | 30.1 | SRRxxxxxxxx |
| MadKes2 | Madagascar kestrel | Falco newtoni | M | 24.0 | SRRxxxxxxxx |
| MadKes3 | Madagascar kestrel | Falco newtoni | M | 29.4 | SRRxxxxxxxx |
| MadKes343 | Madagascar kestrel | Falco newtoni | M | 28.2 | SRRxxxxxxxx |
| MadKes4 | Madagascar kestrel | Falco newtoni | M | 32.7 | SRRxxxxxxxx |
| MK_413 | Mauritius kestrel | Falco punctatus | M | 65.8 | SRRxxxxxxxx |
| MK_421 | Mauritius kestrel | Falco punctatus | M | 40.2 | SRRxxxxxxxx |
| MK_43 | Mauritius kestrel | Falco punctatus | F | 55.4 | SRRxxxxxxxx |
| MK_446 | Mauritius kestrel | Falco punctatus | M | 44.7 | SRRxxxxxxxx |
| MK_591782 | Mauritius kestrel | Falco punctatus | M | 50.3 | SRRxxxxxxxx |
| MK_591798 | Mauritius kestrel | Falco punctatus | F | 46.0 | SRRxxxxxxxx |
| MK_594332 | Mauritius kestrel | Falco punctatus | F | 40.1 | SRRxxxxxxxx |
| MK_594337 | Mauritius kestrel | Falco punctatus | F | 45.0 | SRRxxxxxxxx |
| MK_594338 | Mauritius kestrel | Falco punctatus | F | 41.5 | SRRxxxxxxxx |
| MK_594341 | Mauritius kestrel | Falco punctatus | M | 49.4 | SRRxxxxxxxx |
| MK_594342 | Mauritius kestrel | Falco punctatus | M | 42.2 | SRRxxxxxxxx |
| MK_594345 | Mauritius kestrel | Falco punctatus | F | 42.8 | SRRxxxxxxxx |
| MK_594350 | Mauritius kestrel | Falco punctatus | F | 41.6 | SRRxxxxxxxx |
| MK_594356 | Mauritius kestrel | Falco punctatus | F | 44.5 | SRRxxxxxxxx |
| MK_594357 | Mauritius kestrel | Falco punctatus | M | 39.1 | SRRxxxxxxxx |
| MK_594358 | Mauritius kestrel | Falco punctatus | M | 40.1 | SRRxxxxxxxx |
| MK_594369 | Mauritius kestrel | Falco punctatus | M | 45.9 | SRRxxxxxxxx |
| MK_594370 | Mauritius kestrel | Falco punctatus | F | 49.4 | SRRxxxxxxxx |
| MK_594371 | Mauritius kestrel | Falco punctatus | F | 42.3 | SRRxxxxxxxx |
| MK_594373 | Mauritius kestrel | Falco punctatus | M | 52.2 | SRRxxxxxxxx |
| MK_594387 | Mauritius kestrel | Falco punctatus | M | 40.6 | SRRxxxxxxxx |
| MK_596254 | Mauritius kestrel | Falco punctatus | F | 44.6 | SRRxxxxxxxx |
| RK1 | South African rock kestrel | Falco rupicolus | M | 23.9 | SRRxxxxxxxx |
| RK10 | South African rock kestrel | Falco rupicolus | M | 18.5 | SRRxxxxxxxx |
| RK2 | South African rock kestrel | Falco rupicolus | F | 18.6 | SRRxxxxxxxx |
| RK3 | South African rock kestrel | Falco rupicolus | F | 19.1 | SRRxxxxxxxx |
| RK4 | South African rock kestrel | Falco rupicolus | M | 22.8 | SRRxxxxxxxx |
| RK5 | South African rock kestrel | Falco rupicolus | M | 18.9 | SRRxxxxxxxx |
| RK6 | South African rock kestrel | Falco rupicolus | F | 20.2 | SRRxxxxxxxx |
| RK7 | South African rock kestrel | Falco rupicolus | F | 21.8 | SRRxxxxxxxx |
| RK8 | South African rock kestrel | Falco rupicolus | M | 19.6 | SRRxxxxxxxx |
| RK9 | South African rock kestrel | Falco rupicolus | F | 18.1 | SRRxxxxxxxx |
| SK_D40625 | Seychelles kestrel | Falco araea | M | 26.9 | SRRxxxxxxxx |
| SK_D40627 | Seychelles kestrel | Falco araea | F | 25.1 | SRRxxxxxxxx |
| SK_D40628 | Seychelles kestrel | Falco araea | F | 27.0 | SRRxxxxxxxx |
| SK_SeyK_5965 | Seychelles kestrel | Falco araea | M | 25.5 | SRRxxxxxxxx |
| SK_SeyK_5966 | Seychelles kestrel | Falco araea | F | 33.2 | SRRxxxxxxxx |
| SK_SeyK_5967 | Seychelles kestrel | Falco araea | F | 27.4 | SRRxxxxxxxx |
| SK_SeyK_5968 | Seychelles kestrel | Falco araea | F | 27.4 | SRRxxxxxxxx |

**Supplementary Table 3.** Statistical models of LRS in relation to measures of genome erosion.

(a) Genome Heterozygosity (**het**)

| **Coefficients** | **Estimate** | **Standard Error** | **Z value** | **P value** |
| --- | --- | --- | --- | --- |
| (Intercept) | 0.6170 | 0.2430 | 2.539 | 0.0111 * |
| breeding years | 0.2673 | 0.0356 | 7.509 | 5.95e-14 *** |
| sex | 0.1111 | 0.2064 | 0.538 | 0.5903 |
| scale(**het**) | 0.5682 | 0.1424 | 3.989 | 6.64e-05 *** |

(b) **F_ROH_**

| **Coefficients** | **Estimate** | **Standard Error** | **Z value** | **P value** |
| --- | --- | --- | --- | --- |
| (Intercept) | 0.46375 | 0.28888 | 1.605 | 0.108414 |
| breeding years | 0.30452 | 0.04148 | 7.342 | 2.1e-13 *** |
| sex | 0.10886 | 0.20117 | 0.541 | 0.588404 |
| scale(**F_ROH_**) | -0.68794 | 0.19934 | -3.451 | 0.000558 *** |

(c) Homozygous deleterious alleles (**SnpEff** LOF mutations)

| **Coefficients** | **Estimate** | **Standard Error** | **Z value** | **P value** |
| --- | --- | --- | --- | --- |
| (Intercept) | 0.65062 | 0.24948 | 2.608 | 0.009109 ** |
| breeding years | 0.27727 | 0.03672 | 7.552 | 4.29e-14 *** |
| sex | 0.15936 | 0.20812 | 0.766 | 0.443829 |
| scale(**LOF**) | -0.42502 | 0.11295 | -3.763 | 0.000168 *** |

(d) Homozygous deleterious alleles (Top 5% **CADD**)

| **Coefficients** | **Estimate** | **Standard Error** | **Z value** | **P value** |
| --- | --- | --- | --- | --- |
| (Intercept) | 0.56393 | 0.31144 | 1.811 | 0.0702 |
| breeding years | 0.31428 | 0.04863 | 6.463 | 1.02E-10 *** |
| sex | 0.1378 | 0.25154 | 0.548 | 0.5838 |
| scale(**CADD**) | -0.43744 | 0.17436 | -2.509 | 0.0121 * |

**Supplementary Table 4**. Parameterisation of mortality and breeding across historical and counterfactual epochs in the Mauritius Kestrel simulation model.

| **Epoch** | **Time period** | **Scenario 0: continued conservation** | **Scenario 1: no conservation ever** | **Scenario 2: conservation stops after 2025** |
| --- | --- | --- | --- | --- |
| **1. Baseline ancestral conditions** | ≤1800 | Mortality: 1.00× baseline; breeding: 0.80 | Mortality: 1.00× baseline; breeding: 0.80 | Mortality: 1.00× baseline; breeding: 0.80 |
| **2. Early degradation** | 1800–1950 | Mortality: U(1.00, 1.10)× baseline; breeding: U(0.70, 0.80) | Mortality: U(1.00, 1.10)× baseline; breeding: U(0.70, 0.80) | Mortality: U(1.00, 1.10)× baseline; breeding: U(0.70, 0.80) |
| **3. Late degradation** | 1950–1970 | Mortality: U(1.10, 1.20)× baseline; breeding: U(0.50, 0.70) | Mortality: U(1.10, 1.20)× baseline; breeding: U(0.50, 0.70) | Mortality: U(1.10, 1.20)× baseline; breeding: U(0.50, 0.70) |
| **4. Crash / bottleneck** | 1970–1974 | Mortality: U(1.68, 1.80)× baseline; breeding: U(0.10, 0.20); enforced bottleneck to 4 individuals | Mortality: U(1.68, 1.80)× baseline; breeding: U(0.10, 0.20); enforced bottleneck to 4 individuals | Mortality: U(1.68, 1.80)× baseline; breeding: U(0.10, 0.20); enforced bottleneck to 4 individuals |
| **5. Intensive management** | 1974–1994 | Mortality: U(0.40, 0.50)× baseline; breeding: 0.75; captive releases included | Mortality: U(1.40, 1.50)× baseline; breeding: U(0.10, 0.20) | Mortality: U(0.40, 0.50)× baseline; breeding: 0.75; captive releases included |
| **6. Supportive management** | 1994–2025 | Mortality: U(0.75, 0.80)× baseline; breeding: U(0.50, 0.70) | Mortality: U(1.40, 1.50)× baseline; breeding: U(0.10, 0.20) | Mortality: U(0.75, 0.80)× baseline; breeding: U(0.50, 0.70) until 2025 |
| **7. Future scenario** | after 2025 | Mortality: U(0.75, 0.80)× baseline; breeding: U(0.50, 0.70) | Mortality: U(1.40, 1.50)× baseline; breeding: U(0.10, 0.20) | Mortality: U(1.10, 1.20)× baseline; breeding: U(0.30, 0.40) |

**Supplementary Table 5**. Sensitivity of simulation-anchored genetic load calibration estimated with leave-one-out cross-validation showing estimates being robust.

(a) Calibration with corrected counts

| **Class** | **Full estimate** | **Leave-one-out range** | **Leave-one-out median** |
| --- | --- | --- | --- |
| HIGH | 0.00953 | 0.00746 – 0.0106 | 0.00956 |
| MODERATE | 0.00141 | 0.00138 – 0.00146 | 0.00141 |
| Top 1% CADD | 0.0130 | 0.0124 – 0.0139 | 0.0130 |
| Top 5% CADD | 0.00150 | 0.00140 – 0.00157 | 0.00151 |

(b) Calibration with raw counts

| **Class** | **Full estimate** | **Leave-one-out range** | **Leave-one-out median** |
| --- | --- | --- | --- |
| HIGH | 0.0152 | 0.0135 – 0.0161 | 0.0152 |
| MODERATE | 0.00119 | 0.00117 – 0.00123 | 0.00119 |
| Top 1% CADD | 0.0141 | 0.0131 – 0.0153 | 0.0141 |
| Top 5% CADD | 0.00128 | 0.00116 – 0.00140 | 0.00128 |
